# Genome-wide sweeps create fundamental ecological units in the human gut microbiome

**DOI:** 10.1101/2024.05.25.595854

**Authors:** Xiaoqian Annie Yu, Cameron R. Strachan, Craig W. Herbold, Michaela Lang, Christoph Gasche, Athanasios Makristathis, Nicola Segata, Shaul Pollak, Adrian Tett, Martin F. Polz

## Abstract

The human gut microbiome is shaped by diverse selective forces originating from the host and associated environmental factors, and in turn profoundly influences health and disease. While the association of microbial lineages with various conditions has been shown at different levels of phylogenetic differentiation, it remains poorly understood to what extent unifying adaptive mechanisms sort microbial lineages into ecologically differentiated populations. Here we show that a pervasive mechanism differentiating bacteria in the microbiome are genome-wide selective sweeps, leading to population structure akin to global epidemics across geographically and ethnically diverse human populations. Such sweeps arise when an adaptation allows a clone to outcompete others within its niche followed by re-diversification, and manifest as clusters of closely related genomes on long branches in phylogenetic trees. This structure is revealed by excluding recombination events that mask the clonal descent of the genomes, and we find that genome-wide sweeps originate under a wide regime of recombination rates in at least 66 taxa from 25 bacterial families. Estimated ages of divergence suggest sweep clusters can spread globally within decades, and this process has occurred repeatedly throughout human history. We show, as an example, that the ecological differentiation of sweep clusters forms populations highly associated with age and colorectal cancer. Our analysis elucidates an evolutionary mechanism for the observation of stably inherited strains with differential associations and provides a theoretical foundation for analyzing adaptation among co-occurring microbial populations.

## Introduction

There is widespread agreement that the microbiome associated with our bodies heavily influences our wellbeing. Evidence for such assertions usually stems from correlations of some unit of bacterial diversity with disease or other dysbiotic states^1–5^. Such hypotheses can subsequently be tested in experimental models using animals colonized with isolates assumed to represent the identified bacterial group^6,7^.

However, because correlations with host phenotypes are based on operationally defined units of microbial diversity, ranging from rRNA-to strain-level variants^1–5,8,9^, it is difficult to ascertain whether these units and their associated model isolates accurately represent the adaptive process underlying the association. A recently proposed path forward is to define populations that are both ecologically and genetically differentiated from their sisters because they are the product of divergent environmental selection^10–13^.

Termed reverse ecology, this approach has the potential to more precisely identify the genetic unit associated with host phenotypes since it is optimized by selection to occupy a particular niche space and thus prevail over closely related competitors. Depending on the relative strength of selection to recombination, populations can be either optimized by genes or alleles sweeping across a population, or by a genome-wide selective sweep (GWSS) where the entire genome hitchhikes with an adaptive mutation^14,15^. The latter process has been postulated to be widespread based on theoretical considerations but evidence for its occurrence in nature remains limited^16–18^. We noted, however, that phylogenies of isolates originating from human microbiomes frequently manifest as “brooms”, i.e., an expansion of closely related strains (broom head) on unexpectedly long branches (broom handle). This structure is a hallmark of GWSSs, and arises when an adaptive mutation allows a strain to outcompete all sisters within their specific niche, followed by diversification of the winner into a cluster of closely related strains^19,20^ (Fig. 1a). These considerations therefore led us to hypothesize that GWSSs are a common adaptive mechanism within the human microbiome.

**Figure 1.**
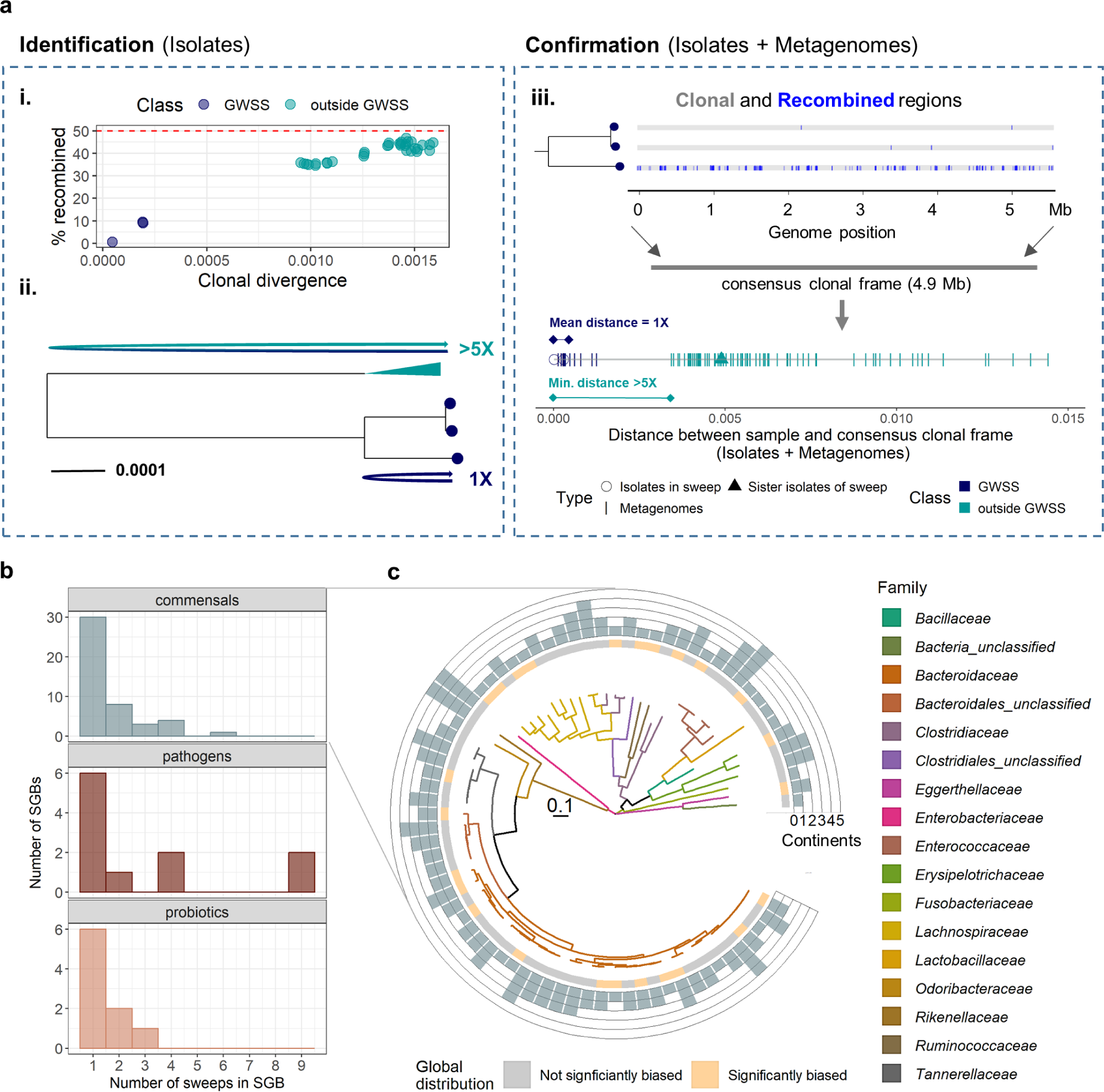
Genome-wide selective sweeps are prevalent in the gut microbiome. **a.** Pipeline used for identification and confirmation of genome-wide sweeps using *Bacteroides intestinalis* (SGB1846) as an example. (i) Identification of GWSS for clusters of genomes that have an average pairwise recombined portion <50% and fulfill the 5X rule (divergence between the cluster and its sister clade exceeds five times the average divergence within the cluster). (ii) Typical “broom-like” structure of GWSS clusters and illustration of the 5X rule. (iii) Confirmation of GWSS clusters by first calculating the consensus clonal frame for each GWSS cluster, then mapping and calculating the distance of all samples (isolate genomes and metagenomes) to the consensus clonal frame, and finally checking adherence to a modified version of the 5X rule (within and between distances are calculated with reference to the clonal frame). The consensus clonal frame consists of the majority nucleotides from concatenated genome regions devoid of recombination in all isolates of a GWSS cluster. **b.** Histogram of the number of GWSS clusters per SGB for commensal, pathogenic, and probiotic bacteria. **c.** Phylogenetic and geographic distribution of GWSS clusters for commensal gut bacteria. The tree is constructed with approximately-maximum-likelihood estimation based on concatenated bac120 marker genes in the GTDB-Tk database (R214, ref. ^43^) for all GWSS clonal frames. Branches of the tree are colored according to bacterial families. The innermost circle represents whether samples in the GWSSs had significantly biased continental occurrence relative to the geographical distribution of other samples in the SGB, based on a Fisher’s exact test (p<0.05); height of the bars on the outer circles indicates the number of continents each sweep covered.

We reasoned that to test the hypothesis that GWSSs create differentiated populations, it is necessary to constrain the effects of horizontal gene transfer and potential isolation bias among available isolate genomes, because both can lead to the observation of long branches in phylogenetic trees^21^. Accordingly, we first sought to identify the vertical descent among closely related genomes by differentiating clonally inherited and recombined portions of the genome, then adopted a theoretical model that estimates the likelihood of genome clusters on long branches arising by GWSSs, and finally mapped metagenomes onto these clusters to test whether intermediate genotypes are missing from isolate collections. We also tested for signatures of selection and differential association with human phenotypes as a proxy for different ecological conditions in the gut and found that GWSS clusters occupy different niche spaces. The intriguing outcome of this analysis is that many human microbiome taxa have been subject to GWSSs, which can spread across the globe over decadal timeframes leading to epidemic-like population structure.

## Results

### Genome-wide sweeps are a common adaptive mechanism

The process of diversification after a GWSS creates the characteristic “broom-like” structures on phylogenetic trees of isolates due to the elimination of genomic diversity within a niche followed by its re-diversification, but the confident identification of such sweep clusters requires that the influence of recombination and isolation bias be constrained. Because homologous recombination can obscure clusters but also generate long branches, especially among closely related genomes^21^, we first sought to identify the clonally inherited portion among genome pairs. To this end, we built on previous approaches that partition the SNP distribution between pairs of genomes into a mixture of a Poisson and a negative binomial distribution^10,22^. While the former captures the accumulation of point mutations in clonal genome regions, the latter models the effect of recombination, which can either elevate or reduce SNP density depending on the phylogenetic distance of the source genome (Extended Data Fig. 1, Methods).

By determining a best fit for the observed SNP distribution to a mixture of these two models, we can estimate (i) the fraction of the genome that is vertically inherited versus recombined, (ii) the divergence in each fraction, as well as (iii) the recombination rates between pairs of genomes. We comprehensively assessed this approach with simulated sets of genomes evolved from a single ancestral genome while incorporating selection and recombination under a wide range of mutation and recombination rates, and defined the regime where recombination can be confidently differentiated from point mutations (Extended Data Fig. 2, Methods). Because this divergence regime was found to be at the subspecies level, we delineated species-level genome bins (SGBs) as groups of genomes that are <6% diverged from a reference genome in the MetaPhlAn4 database^23^ and applied the mixture model to each SGB separately. We used a dataset of 16,864 human gut isolate genomes, which we dereplicated according to human subjects to avoid double counting of clonal isolates from the same individual due to repeated sampling (Methods). This categorization resulted in 176 SGBs containing >5 genomes, which showed widely varying fractions of clonal and recombined genome portions, suggesting that the diversity within SGBs is differentially structured by vertical and horizontal inheritance (Extended Data Fig. 3).

To test for the occurrence of GWSSs, we first analyzed the isolate genomes comprising the 176 SGBs and subsequently used metagenomes to confirm the observed structure to account for potentially missing diversity from the isolate collections. We developed PopCoGenomeS (Populations as Clusters of Genome Sweeps), a computational pipeline to search the 176 SGBs for groups of genomes that were vertically inherited from a common ancestor and displayed hallmarks of GWSSs. We tested whether there is support for GWSSs by applying a coalescent-based model (5X rule) that compares the diversity within a cluster of genomes to its closest relative (Fig. 1a)^24,25^. If the ratio of between to within cluster divergence is >5, the probability that an identified GWSS is in fact due to evolutionary drift, the most likely alternative mechanism to selection for generation of sequence clusters, is <1% (Methods). In this way, we identified 377 putative GWSS clusters with many SGBs containing multiple clusters, and asked whether this structure can be confirmed by whole community metagenomics that is not subject to isolation bias by compiling 1477 metagenomes from 74 datasets covering different age, disease state, and biogeography (Methods). We first determine a majority-rule consensus sequence for each GWSS cluster, which is the majority nucleotide in each position of the genome not affected by recombination (consensus clonal frame) (Fig. 1a). Then, we calculated the distance of each of the 1477 metagenomes as well as all isolate samples to this consensus clonal frame for the confirmation of GWSS clusters that adhere to the 5X rule (Fig. 1a, Methods).

In this combined isolate genome and metagenomic dataset, we were able to confirm 124 GWSSs in 66 SGBs, which consisted of 46 commensals, 11 pathogens, as well as 9 commensals that are frequently found in fermented and functional foods (probiotics) (Fig. 1b, Tables S1 and S2, see Methods for SGB classification). Although the majority of SGBs contained a single GWSS cluster, many contained multiple. This indicates that species operationally defined by average nucleotide identity can consist of several evolutionarily defined units of diversity, which, based on the evidence presented further below, display differential ecology. Specifically focusing on commensals where the identification of sweeps is less likely to be affected by disease outbreaks or the widespread use of probiotics, we found a total of 77 GWSSs in 46 SGBs spanning 17 bacterial families. While *Bacteroidaceae* had the highest prevalence of GWSS clusters, they are found in most of the other major human gut bacterial families such as *Clostridiaceae, Lachnospiraceae and Enterococcaceae* indicating that GWSSs are a pervasive mechanism differentiating populations in the human microbiome (Fig. 1c). Since the number of sweeps detected in each SGB significantly correlates with the total number of samples where the SGB is detectable (Spearman’s test, n=46, Rho=0.46, p=0.001), it is likely that GWSSs exist in other gut bacterial families that have been less extensively cultured.

Further support for GWSS clusters being selectionally optimized units of diversity arises from population genetic metrics. Under population genetic theory, Tajima’s D quantifies the overrepresentation of rare alleles in a population and a negative value indicates that it is recovering from a selective sweep or a population bottleneck^26^. Indeed, all commensal GWSSs we observed had negative Tajima’s D values (Extended Data Fig. 4a). Moreover, more than half of the sweep clusters (43/77) contained at least one gene that showed signatures of very strong positive selection as indicated by the ratio of non-synonymous to synonymous substitutions (dN/dS) being larger than 2 (Extended Data Fig. 4b)^27,28^. Taken together, these measures are consistent with positive selection playing a role in diversifying microbiome bacteria by GWSSs and led us to investigate how widespread GWSSs are geographically as further evidence for their adaptive nature.

Although extreme and persistent bottlenecks could lead to the erroneous inference of GWSS, the likelihood of this scenario becomes diminishingly small if GWSS clusters are widely distributed among human populations. Indeed, the majority of GWSS clusters (54/77) comprise isolates or metagenomes from more than one continent, indicating that they have spread among different human populations (Fig. 1c). Among these 54 widespread clusters, 42 show no spatial preference (Fisher’s exact test, p>0.05 for the geographical distribution of samples inside and outside a GWSS cluster, Fig. 1c), suggesting that they respond to similar selective pressures in diverse human populations. Remarkably, one *Bacteroides uniformis* sweep cluster was unique to indigenous populations, being shared among three populations on two different continents: the Baka and Beti people in Cameroon^29^, as well as the Matses people in Peru^30^. Because the Baka and the Matses are geographically remote hunter-gatherer communities that have never been in direct contact with each other, the spread of the same sweep cluster among them is best explained by a chain of transmission events mediated by less isolated populations. Moreover, the absence of this GWSS cluster from other populations indicates that the transmission occurred in the past, with the relevant strains now extinct in most human populations, possibly due to lifestyle shifts related to industrialization and urbanization. These observations of global distributions of GWSS clusters lead us to ask on what approximate time scales sweeps may occur, especially since fast global spread would further support that GWSS clusters are the product of selection rather than population bottlenecks and drift.

### Sweep clusters arose repeatedly and spread rapidly

We find that GWSSs have happened repeatedly over the course of human history by constraining the age of sweep clusters using two independent methods. First, based on a previously determined molecular clock rate for commensal and pathogenic bacteria of 1-10 mutations per genome per year^31–33^, we estimate that the age of sweep clusters ranges from approximately 10 to 1000s of years (Fig. 2a). These age estimates are consistent with those obtained by a second method that independently estimates molecular clocks by determining the number of substitutions that have accumulated within the same GWSS clusters detected in metagenomes of individuals who were sampled at least twice with a time interval of >1 year, or twins who have resided apart since reaching adulthood (Methods). With only two exceptions, the age of sweep clusters estimated from these metagenome-based evolutionary clocks was within the same order of magnitude to those estimated using the fixed molecular clocks of 1-10 SNPs genome^−1^ yr^−1^ (Fig. 2b). Although we realize that these estimates carry large uncertainties, they still suggest that GWSSs have happened repeatedly over the past thousands of years with clusters showing nearly a 1,000-fold difference in age, and that even within a taxonomic species (SGB), sweep clusters with orders of magnitude different ages can exist (Fig. 2a).

**Figure 2.**
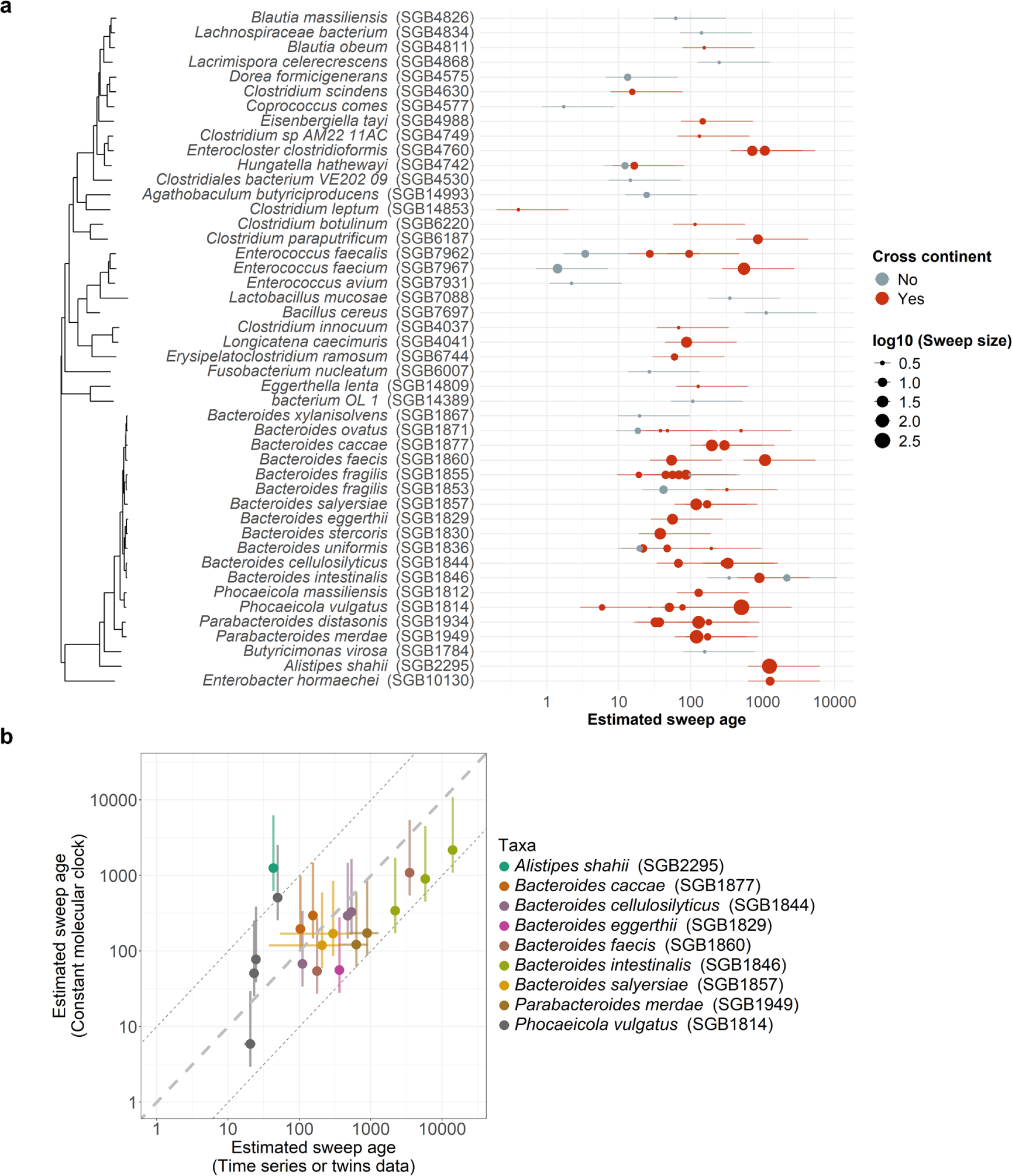
Genome-wide selective sweeps arose repeatedly and spread rapidly. **a.** Estimated age of GWSS clusters (dots) using a constant molecular clock with a median mutation rate of 5 SNPs genome^-1^ year^-1^, with the size of the dot representing the number of samples (isolate genomes and metagenomes) in each GWSS cluster. Error bar for each dot represents the upper and lower limit of the age of each sweep cluster at 1 and 10 SNPs genome^-1^ year^-1^, respectively. The color of the dots indicates whether a sweep cluster was detected across more than one continent. The approximately-maximum-likelihood tree on the left is based on concatenated bac120 marker genes (GTDB-Tk database, R214, ref. ^43^) extracted from a randomly selected GWSS clonal frame in each SGB. **b.** Comparison of estimated ages of sweep clusters using a constant molecular clock (5 SNPs genome^-1^ year^-1^) and metagenomic molecular clocks determined separately for each SGB. The gray dotted line in the middle represents a 1:1 ratio between the two estimates, while the two gray dotted lines on the sides represent 10:1 and 1:10 ratios, respectively. Horizontal error bars represent 95% confidence intervals of the estimated age from metagenomes, while vertical error bars represent the upper and lower limits of the sweep age under a constant molecular clock (1 and 10 SNPs genome^-1^ year^-1^).

Since the age of sweep clusters also provides an upper bound for the time to spread into geographically distant human populations, we asked what is the minimum time we can estimate from the data for transmission across continents. Using the median molecular clock of 5 SNPs genome^−1^ yr^−1^, a total of 26 multi-continental sweep clusters were estimated to be younger than a century (Fig. 2a). Furthermore, we found that in the majority (47/54) of multi-continental sweep clusters, the genetic distances of samples originating from different continents were not significantly different from those from the same continent (ANOSIM test, p>0.1). This result indicates that in most SGBs, there were rapid and repeated cross-continental transfers of strains. Interestingly, the cross-continental *B. uniformis* sweep unique to the three indigenous and reputedly isolated populations mentioned above was the oldest GWSS in *B. uniformis*, with an estimated age of approximately 100-1000 years, supporting that this cluster may have spread before industrialization. Finally, the average SNP diversity within sweeps for commensal bacteria is significantly larger than that for pathogenic bacteria (Student t-test on log-transformed data, n=77 and 34, df=63, p=0.018, Extended Data Fig. 5). Therefore, the transmission of commensal bacteria within the human population is slower and /or the sweep clusters are older compared to infectious pathogen outbreaks. Nonetheless, our data indicate that even commensal GWSS clusters can spread globally within the lifespan of a human and their distribution is consistent with shared selective regimes in diverse human hosts.

### GWSSs happen across a wide range of recombination rates

Theory predicts that since GWSSs occur when the entire genome hitchhikes with an adaptive mutation, the relative rate at which the genome carrying the adaptation spreads within its niche must be much higher than that of the adaptive gene or allele being shared by recombination^14,15^. This is because high recombination rates can lead to gene-specific sweeps by breaking the linkage between the beneficial mutation and its neighboring variants, allowing the adaptive allele to spread independently of its genomic background. Contrary to this explicit expectation, we find that SGBs with no confirmed GWSSs have comparable recombination rates (genome fraction recombined per mutation) to those containing GWSSs (median 2.4×10^-4^ vs. 1.7×10^-4^; Wilcoxon rank sum test, n=52 and 45, p=0.09; Fig. 3, Extended Data Fig. 6). In fact, among commensals, GWSSs were detected across a range of recombination rates that varied approximately 400-fold. For example, *Blautia massiliensis* (SGB4826) harbors a sweep despite being among the fastest recombining SGBs, while multiple other SGBs from the same genus have recombination rates that are 3-20 fold slower with no observed sweeps (Fig. 3). Our data thus suggest that selection is frequently strong enough to permit GWSSs even under high recombination regimes.

**Figure 3.**
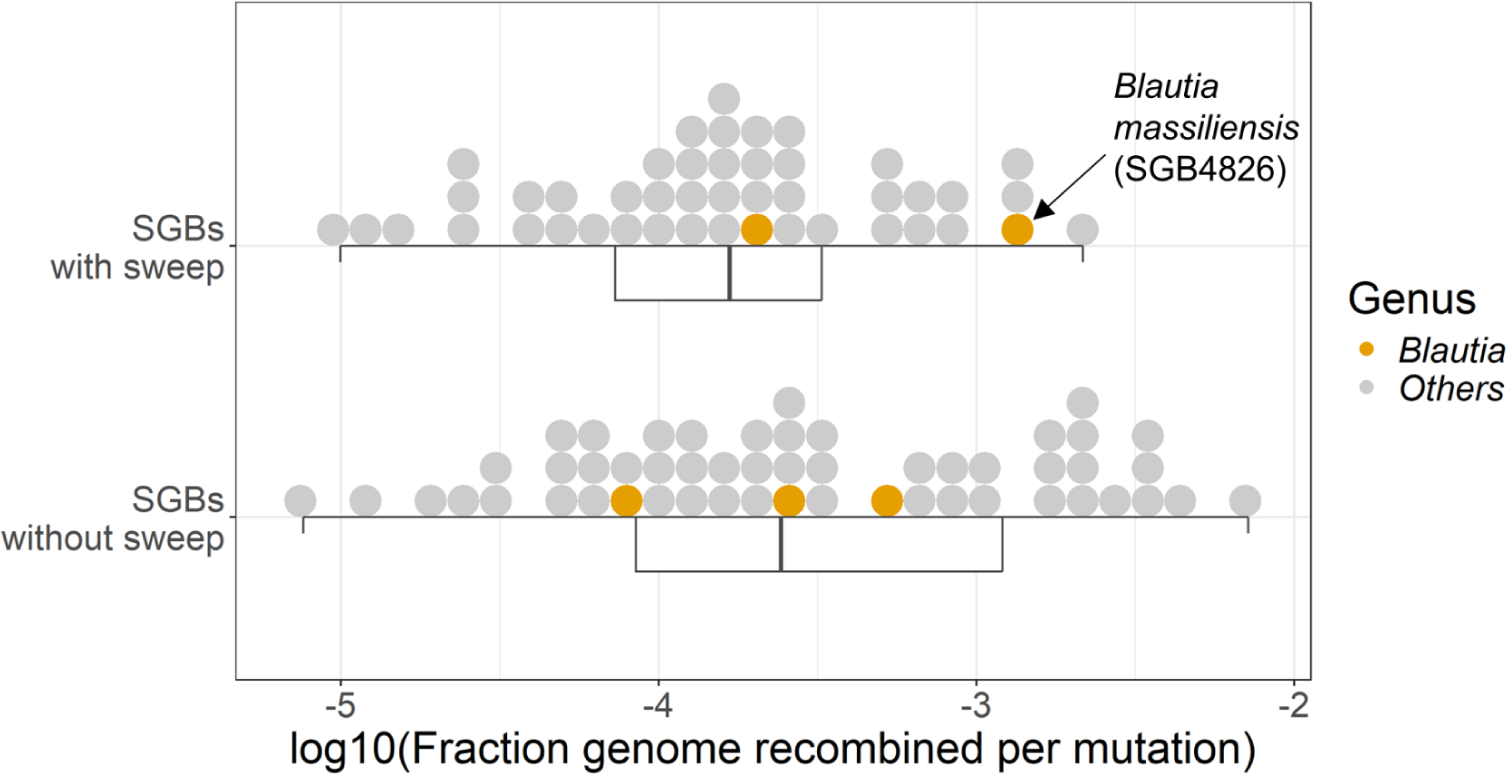
Genome-wide selective sweeps occur across a wide range of recombination rates. Comparison of the recombination rates between SGBs with (top panel) and without (bottom panel) detected GWSSs, with each dot representing an SGB. The box plot below each dot plot indicates the interquartile range (IQR, box), median (middle line), and the most extreme data points within 1.5×IQR of the upper or lower quartile (whiskers) of the distribution.

### GWSS clusters represent ecologically differentiated populations

A key prediction of the theory underlying the occurrence of GWSSs is that sweep clusters are ecologically differentiated. Although fine-scale niche differences in the gut are difficult to constrain, we hypothesized that major life-history or physiological changes in humans may lead to shifts in ecological conditions that bacterial populations may respond to. As a test case, we searched for associations of GWSS clusters with two phenotypes, colorectal cancer (CRC) and seniority due to their ample representation in datasets across different human populations. We extended the 5X rule to phylogenetic distances based on alignment of metagenomes to marker genes in the MetaPhlAn database as this approach allows more efficient scaling up to large datasets (Methods, Extended Data Fig. 7). We were thus able to expand the screening for GWSS clusters to a collection of 3,941 metagenomes from 17 datasets covering various geographies. Using a general linear model with a forward, stepwise variable selection approach, we found 30 and 27 GWSS clusters with significant associations with CRC and seniority, that are FDR corrected at 0.05 and span across multiple datasets with no significant geographical bias (chi-squared test within each GWSS, p>0.05, Extended Data Fig. 8). Using the same approach to analyze datasets within countries with CRC subjects, we found 28 and 26 GWSSs that had country-specific associations with CRC and seniority, respectively.

In line with GWSS clusters being finely differentiated ecological units, clusters within the same SGB can have opposing associations with the same host state (Fig. 4). In 5 SGBs such opposing associations were detected in the combined dataset of all countries with no significant bias for any one country, while an additional 5 SGBs contained GWSS with opposing associations within one or more countries (Extended Data Fig. 8). Most of the SGBs (4 of 5) with bidirectional cross-country associations were *Bacteroidales*. These SGBs generally contained high numbers of sweep clusters, each with limited prevalence, suggesting fine-scale and frequent differentiation within this order (Fig. 2a). For example, while *Bacteroides fragilis* (SGB1855) was not significantly associated with CRC or seniority (Methods, Fig. 4, Extended Data Fig. 8), its GWSS clusters were associated with both host phenotypes in different directions (positive to negative: 3:2 in seniority and 1:1 in CRC). It was also one of the SGBs with the highest numbers of confirmed sweeps (6 GWSS clusters, Fig. 2a), with the average sweep cluster comprising only 7.2±3.9 samples. This indicates contrasting selection within the same taxonomic species and is consistent with the ability of strains with a selective advantage to quickly occupy a niche, the range of which constrains the extent of occurrence of the sweep cluster. To explore potential functions underlying such niche differentiation, we used *B. fragilis* (SGB1855) as a model and asked what gene clusters (i.e., contiguous genes that likely represent operons) might be specific to each GWSS cluster (Methods). This analysis revealed enrichment in functions involved in the synthesis or modification of glycans and glycoconjugates, such as capsular polysaccharides (Extended Data Fig. 9, Table S3). As these structures have long been known to contribute to pathogenicity by allowing pathogens to escape specific immune responses^34–36^, it is possible that commensal bacteria use similar mechanisms to adapt to discrepant host phenotypes related to health or disease. Indeed, *B. fragilis* capsular polysaccharides can protect against inflammatory disease^37^, and genes regulating capsular biosynthesis have been inferred to be under adaptive evolution within individual human hosts^32^. Altogether our analysis suggests that GWSS clusters in commensal bacteria can display highly specific associations with human phenotypes, and their population structure may be shaped by interaction with the immune system.

**Figure 4.**
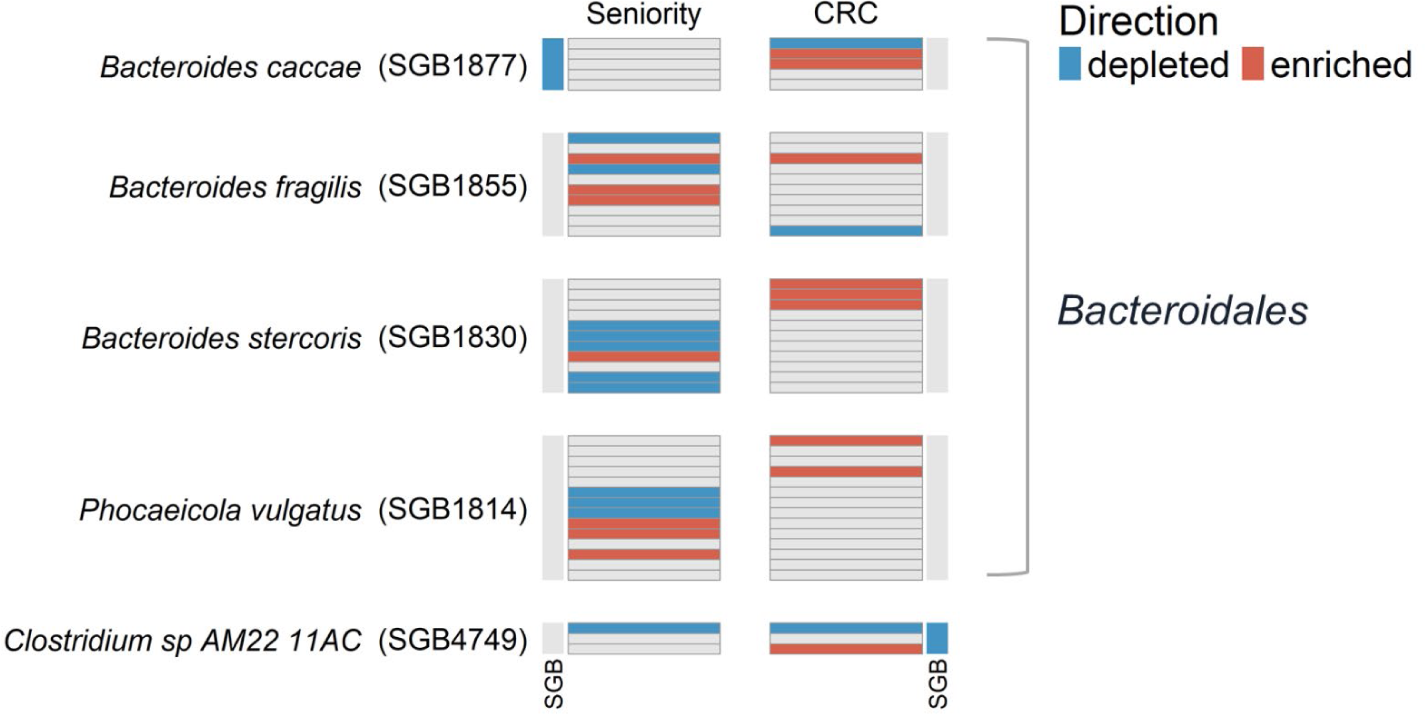
Contrasting ecological associations of genome-wide selective sweep clusters within SGBs. Heatmaps of statistically significant (p_adj_<0.05) associations between GWSS clusters and seniority (left) or CRC (right). All associations shown are those that span multiple datasets and show no significant geographical bias (p>0.05, chi-squared test). Vertical bars on the side of each panel represent whether the SGB that the GWSS clusters belong to is significantly associated with seniority or CRC.

## Discussion

Our analysis suggests that GWSSs are a fundamental force structuring bacterial populations within humans and we suspect that they are even more common since our analysis was limited by the availability of sets of closely related isolate genomes. Because GWSS create ecologically differentiated populations, their mapping onto metagenomes may serve as accurate and easily definable markers for a variety of conditions. We chose CRC and seniority as test cases, but it will be interesting to expand mapping of GWSS clusters to other human phenotypes, especially to test if a particular cluster shows multiple associations, as this may point to shared properties among disease or dysbiotic states. A further important corollary of their ecological differentiation is that GWSS clusters should be adapted to coexistence with at least a subset of other community members, so that the identification of GWSS clusters may facilitate the construction of bacterial co-occurrence networks, and further impact the choice of appropriate isolates to serve as experimental models or the assembly of synthetic gut communities.

A surprising outcome of our analysis is that GWSS clusters of gut commensals appear globally adaptive and spread in a fashion that resembles epidemics, which are typically ascribed to pathogens^38,39^. However, the underlying mechanisms involved in the spread of commensal and pathogenic bacteria are likely different. While it is known that for many pathogens, novel adaptive clones can spread rapidly, leading to genetically highly homogeneous global populations, our data suggest that commensals may require decades to reach global distribution likely because of slower transmission chains. This slower spread is reflected in higher diversity within commensal GWSS clusters since they may diversify as they spread geographically. Strains, which are often regarded as the lowest identifiable unit of diversity within metagenomes, have been shown to be primarily acquired from related individuals^40^. Strain sharing among unrelated individuals has been observed at both local and global scales^40–42^, but the eco-evolutionary mechanism behind this phenomenon remains unclear, despite suggestions that host adaptation plays an important role^41^. In the model proposed here, although not all closely related strains among unrelated individuals in an SGB originate from the same clonal expansion^42^, a portion can be part of GWSS clusters, and their acquisition reflects primarily their superior adaptation to the specific settings within a microbiome. Indeed, once acquired, strains have been shown to persist over extended periods of time^5^, on average over a third of the human lifespan^40^, and we suggest that their replacement also reflects the turnover of ecologically adapted populations, which may also serve as a sentinel for shifts in human health.

## Supporting information

All Supplementary Figures

## Acknowledgements

We thank the Life Science Compute Cluster at the University of Vienna for providing data infrastructure and access to high-performance computing clusters. We would like to acknowledge the Joint Microbiome Facility of the Medical University of Vienna and the University of Vienna for their support with sample preparation, sequencing and sequence data processing. We thank David Berry and Jessie Shapiro for comments on the manuscript. We also thank Philip Arevalo, Otto Cordero, Jeff Gore, Fangqiong Ling, Stephanie Schnorr, Anni Zhang, and Shijie Zhao for discussions and comments at various stages of the study, as well as Mathieu Groussin and Mathilde Poyet for providing us early access of sequencing data. This study was supported by the Austrian Science Fund (FWF) [doi.org/10.55776/COE7], Vienna Science and Technology Fund (LS18-053) and the Austrian Science Fund (KLI 557). C.R.S. was partially supported by a Fellowship from the Natural Science and Engineering Council of Canada Postgraduate Scholarship-Doctoral (NSERC PGS-D).

## Author contributions

X.A.Y., C.R.S. and M.F.P. developed the concept and designed the study. X.A.Y. and A.T. performed data collection. X.A.Y. and C.R.S. conducted all data analyses. M.L., C.G. and A.M. performed all colonoscopy and collection of the newly sequenced Austrian isolates. X.A.Y. and M.F.P. prepared the manuscript with input from all coauthors.

## Competing interests

The authors declare no conflict of interest.

## Methods

### Isolate genome collection

A total of 19,837 publicly available isolate genomes were collected and consisted of the entire data in the Unified Human Gastrointestinal Genome (UHGG) catalogue^44^ (Version 1.0, 10,648 genomes) and four large-scale culturomics studies^29,45–47^. In addition, a group of 186 isolates from Austrian individuals were newly collected and sequenced. A total of 50 patients undergoing colorectal cancer screening colonoscopy at the Vienna General Hospital were enrolled in the study, comprising 24 individuals with irritable bowel syndrome, 5 with ulcerative colitis, and 21 healthy controls. No subjects were found to have carcinomas during the colonoscopy.

For bacterial isolation in the Austrian study, brush samples and biopsies were collected during colonoscopy from the ileal or cecal mucosa or ascending colon with or without an endoscopically visible biofilm and were processed immediately^48^. Brushes and biopsies were vortexed or homogenized in 0.6 ml of 0.9% NaCl, and the suspensions were subsequently plated on one of the following six culturing conditions: Columbia agar with 5% sheep blood, MacConkey agar, Columbia CNA agar with 5% sheep blood, and CPS agar (Becton Dickinson) under aerobic conditions at 37°C; Brucella agar with 5% horse blood, and Schaedler KV Agar with 5% sheep blood (Becton Dickinson) under anaerobic conditions at 37°C. Aerobic cultures were assessed after 18h and 48h, and anaerobic cultures after 48h and 72h. Colonies identified as *Bacteroides* and *Parabacteroides* by matrix-assisted laser desorption ionization time-of-flight mass spectrometry (MALDI-TOF MS) analysis on a MALDI Biotyper MBT smart instrument (Bruker), or by 16S rRNA gene sequencing on a capillary sequencer (SeqStudio Genetic Analyzer, Applied Biosystems by Thermo Fischer Scientific) were saved as glycerol stocks at −80°C.

Glycerol stocks of *Bacteroides* and *Parabacteroides* were cultured in brain heart infusion medium with supplements for 24 h before DNA isolation. DNA isolation was performed on the King Fisher Flex instrument (Thermo Fisher Scientific) using the MagMAX™ DNA Multi-Sample Ultra 2.0 Kit (Thermo Fisher Scientific) including an initial proteinase K digestion step and an RNase treatment step.

Sequencing libraries were prepared using the NEBNext® Ultra™ II FS DNA Library Prep Kit, with the NEBNext® Multiplex Oligos for Illumina barcodes. Sequencing was performed on the Illumina NovaSeq 6000 platform using SP flow cells (300 cycles, 2×150bp paired-end). Reads were trimmed, filtered and merged with BBMap 38.90 (ref. ^49^, ktrim=r k=21 mink=11 hdist=2 minlen=125 qtrim=r trimq=15), and *de novo* genome assembly was performed via Spades v3.15.5 (ref. ^50^) under isolate mode.

### Isolate quality filtering and taxonomic assignment

The total of 20,023 genomes collected were evaluated with CheckM v1.2.2 (ref. ^51^) to screen for genomes that met the following criteria: >85% genome completeness, <5% contamination, and N50 >50K. This selection process yielded a total of 16,864 genomes that were of sufficient quality for downstream analysis. We assigned each genome to a species-level genome bin (SGB) according to the MetaPhlAn4 reference genome database (version Jan. 2022, ref. ^23^). Each genome was assigned to the SGB it was most closely related to based on FastANI 1.33 (ref. ^52^) with the centroid genome as the primary reference or, if unavailable, a representative genome. The SGB assignments were only made for genomes whose closest relative had an average nucleotide identity (ANI) >94% and over 30% sequence alignment. To address potential mis-assignments of genomes associated with species possessing species boundaries slightly lower than the most commonly used threshold (95%)^53^, we adopted a lower-end species boundary of 94% ANI for SGB assignment, as opposed to the more commonly used 95% ANI. Genomes failing to meet the 94% ANI cutoff with their closest relatives were converted to synthetic fastq reads (ART-2016.06.05, -ss HS25, ref. ^54^) and assigned by MetaPhlAn4 (version 4.0.3, ref. ^23^). Genomes that could not be assigned by either method were excluded. We further checked if the ANI-based and MetaPhlAn4-based SGB assignments are generally consistent by converting a random subset of genomes in each ANI-based SGB to synthetic fastq reads (ART-2016.06.05, -ss HS25, ref. ^54^) and assigning them by MetaPhlAn4. Although the majority of ANI-based and MetaPhlAn4-based SGB assignments were congruent, certain ANI-based SGBs had all genomes assigned to another SGB in MetaPhlAn4. All genomes in these SGBs were reassigned to their corresponding MetaPhlAn4 assigned SGB (Table S4).

### Isolate genome filtering based on metadata

A key step to ensure unbiased genome-wide sweep identification is to filter the genomes so each SGB only contains isolates originating from different individuals. For each isolate, we retrieved information on human subject identifiers, age, sex, health status, country, year, and creator of collection, and BioProject accession number from the UHGG database^44^, as well as from the text or supplementary materials of the respective publications. For each SGB, we only retained isolates that originated from a different human subject, based on either a unique subject identifier, country of sample, or BioProject number (representing a different study; studies with multiple BioProject numbers were checked for and corrected manually).

For studies with more than 10 genomes but no human subject identifier or country information, we created a human subject ID as a combination of the following five factors: age (or age group when exact age is not available), sex, health status of the subject, year, and creator of collection. Human subjects with different combination-based subject IDs were considered as different individuals. When multiple genomes from a single SGB were isolated from the same individual, we choose the genome with the highest quality score according to dRep v 3.4.1 (ref. ^55^). This procedure resulted in a final collection of 6,411 high-quality isolate genomes that originate from different human subjects (Table S5). These isolates span 995 SGBs, of which 176 contain more than 5 genomes. Since one SGB, SGB10068, assigned as *E. coli*, contained 1,053 genomes and was vastly larger than any other SGB, we randomly subsampled this SGB to 25% of its original size, such that it became comparable in size to the second-largest SGB. This SGB was renamed as SGB10068s to indicate the subsampling. Each SGB was assigned to its corresponding family, genus, and species-level taxonomy in the MetaPhlAn4 database.

### Estimation of recombined and clonal genome fractions via mixture modeling

To estimate the recombined and clonal fractions in a pair of genomes, we developed a method that employs a combination of Maximum Likelihood Estimations (MLE) and Hidden Markov Models (HMM). This method is conceptually similar to that in Sakoparnig *et al.* 2021 (ref. ^22^), but with several technical adjustments (detailed at the end of the model validation section below). The major rationale behind the method is that SNPs introduced by mutations between a pair of genomes should be randomly distributed across the genomes, while recombination generates regions in the genomes that have an increased or decreased number of SNPs depending on whether the recombined genome fragment stems from a distant or close relative (Extended Data Fig. 1). It is therefore possible to partition genome alignments into regions that have been vertically inherited or recombined based on SNP distributions across the alignment.

The SNP distribution for each pair of strains was determined by sliding 500bp windows across pairwise genome alignments with a step size of 50, resulting in a probability mass function P(x=n) of 500bp windows that have n SNPs. This probability mass function was partitioned as a fractional sum of a Poisson distribution that represents the clonal fraction of the genome, and a negative binomial distribution that represents the recombined fraction of the genome using the following equation:

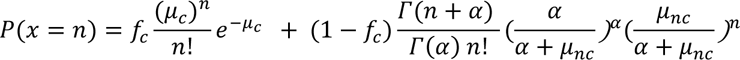

where μ_*c*_ and μ_*nc*_are the means for the SNPs/window in the clonal fraction (Poisson distribution) and the recombined fraction (negative binomial distribution), and α is the dispersion parameter of the negative binomial distribution. The fraction of the genome that is clonally inherited or recombined is represented by *f*_*c*_ and 1 − *f*_*c*_, respectively. The observed SNP distribution was fitted to the equation using maximum likelihood estimation with the L-BFGS-B algorithm in the python package SciPy (ref. ^56^). Since the negative binomial distribution can be interpreted as a Gamma mixture of Poisson distributions with the dispersion parameter α, and approaches the Poisson distribution when α =1, the lower bound of α was set as 2 so that the distribution is sufficiently distinct from the Poisson distribution that represents the clonal fraction. The effective recombination rate r/m, which is the number of SNPs exchanged by recombination relative to the number of SNPs introduced by mutation, can be calculated as μ_n_ (1 − *f*_*cc*_)/ μ_*c*_ *f*_*c*_.

The estimated recombination and clonal fractions are further validated through the utilization of an HMM with the python package pomegranate (ref. ^57^), where the two fractions serve as hidden states (C: clonal state; R: recombined state). The Viterbi training algorithm was applied to the spatial SNP profiles for each pairwise alignment, with the *f*_*c*_and 1 − *f*_*c*_from the MLE as the starting proportion of C:R. Similarly, the initial parameters for the HMM emission matrix were generated from the MLE estimated Poisson and negative binomial distributions. While in most cases the HMM yielded results that were highly consistent with the MLE, the HMM validation step was effective in correcting occasional MLE failures at low clonal divergences. Furthermore, the relative occurrence rates of recombination to mutation (*ρ/θ*) could be estimated as the total number of R states divided by the total number of SNPs in the C state.

### Validation of model performance with simulated data

To assess the performance of our mixture model across the expected biological ranges of evolutionary processes in gut bacteria, we evaluated it on sets of simulated genomes covering a wide range of population genomic parameters. This approach also allowed us to optimize our model so it would be most effective in the divergence range where recombination is expected to influence the detection of GWSSs.

We generated sets of genomes (N=64) with the program CoreSimul (ref. ^58^), a forward simulation program that evolves a single genome along a phylogenetic tree to generate derived genomes while incorporating recombination. For each phylogenetic tree, 144 parameter combinations were tested: (i) the scale (i.e., maximum pairwise distance) of the tree s=0.0002, 0.001, 0.005, 0.02, 0.036, and 0.05, (ii) the size of the recombination fragment, exponential distributions with mean delta=200, 500, 1000, (iii) the relative occurrence rates of recombination to mutation *ρ/θ*=0.01,0.1,0.2,1, and (iv) the rate of exponential decay with divergence for success of recombination Φ=9,18, when p(success)=10^-πΦ^. We simulated the evolutionary process of a 2 million base pair genome diverging into 64-genome populations with 2 different types of phylogenetic structures: one where multiple genome-wide sweeps have occurred (Extended Data Fig. 2a), and another one where the tree is fully balanced (Extended Data Fig. 2g). During each time segment (i.e., time between consecutive nodes) on the tree, each branch on the tree receives mutations (Jukes-Cantor 69 substitution model) and recombination events based on a Poisson process, but only branches that overlap in time are allowed to recombine with each other, and the probability of successful recombination exponentially decreases with sequence divergence using empirically observed rates^59^. For each pair of genomes, we track all regions that have undergone recombination since their last recent common ancestor, and regions with overlapping recombination events are merged and treated as a single event. Finally, we applied our mixture model to the simulated genomes and conducted a comparison between the estimated recombined fraction of the genome, the clonal divergence, and two measures of recombination (relative effect of recombination and mutation r/m, and relative recombination to mutation occurrence rate *ρ/θ*) with their corresponding values in the simulation.

We found our method performed well when there were >1,500 total SNPs (>0.075% divergence, including SNPs introduced by both recombination and mutation) in the pairwise alignment, the majority of recombination fragments were >500bps, and the overall recombined fraction of the genome was <2/3 of the genome. We interpret the limitations as follows: First, the method becomes noisy when the overall number of SNPs falls under 1,500, which is likely due to a lack of sufficient SNP-containing windows for either the MLE or the HMM to perform efficiently. Second, our method considerably overestimates the recombined fraction of genomes when the length distribution has a mean of 200, because too many windows (500bp) are only partially recombined, and the fraction of windows that are identified as recombined no longer equals the fraction of the genome that is simulated as recombined. Third, the method also loses accuracy when the mean divergence of the recombined fraction is less than 2.5 times than that of the clonal fraction since the SNP distributions in these two fractions overlap too much to be sufficiently resolved from each other. This mostly occurs when >2/3 of the genome is recombined under our parameter settings.

Considering the above results, we optimized our method so it would perform best in a range characterized by low-to-intermediate levels of genome recombination as expected in GWSS clusters that are of relatively recent origin and hence still retain a high fraction of vertically inherited genome (clonal frame). Our optimization strategy involves using an intermediate window size for counting SNPs, and filtering at both ends of the recombination spectrum where either a very small or very large fraction of the genome was recombined. We opted for a window size of 500bps since the recombination fragment size in bacteria is estimated to range from tens to thousands of base pairs^22,60,61^, and that further using a smaller window size could compromise the resolution of the method at low divergences due to insufficient numbers of SNPs per window. Validation of our method using simulated data allowed us to establish robust filters to ensure accurate parameter estimation, free from the influence of degenerate parameter sets resulting from the MLE being confined to a local minimum. These filters were set for genome pairs that were expected to be very highly or lowly recombined: All genome pairs with less than 1,500 SNPs were considered 100% clonal, with the divergence of the clonal fraction deemed as 10^-5^; meanwhile, all genome pairs for which the estimated mean of the recombined fraction is less than 2.5 times that of the clonal fraction were considered as 100% recombined, with both the clonal and recombined fractions of the genome sharing the same divergence as the overall genome alignment.

Moreover, as a precaution against sporadic failures of the MLE, we implemented a corrective measure. We cross-checked whether the MLE-estimated recombination fraction exceeded twice that determined by the HMM. If such a discrepancy occurred, we substituted the MLE-estimated parameters with those derived from the HMM. Conversely, if no such discrepancy was observed, the MLE-derived clonal divergence and recombination fractions were deemed the final estimated parameters. Since the HMM was also additionally used to determine the spatial information of the recombined regions, we find that most recombined regions that stretch for less than 8 consecutive sliding windows in the HMM are falsely identified, and therefore reassign them as clonal regions after completion of the HMM.

### Identifying putative GWSSs from the isolate collection

We established two criteria for the conservative identification of putative GWSSs in the isolate genome collection, and encapsulate the relevant workflow into a package called PopCoGenomeS (Populations as Clusters of Genome Sweeps). Firstly, to ensure a sufficiently large clonal frame for confident phylogenetic analysis and downstream metagenomic mapping, we only consider genomes that are predominantly vertically inherited (i.e., the pairwise recombined portion is <50%). Secondly, the divergence among the genomes considered should satisfy the 5X rule, which is a stricter variant of the previously established 4X rule^24^. According to the 4X rule, if sister clades on a tree with the same sample size are separated by a distance gap that exceeds 4 times the within-clade distance, there is less than 5% probability that the clades are formed due to random drift. The 5X rule decreases the probability of drift to less than 1% and also allows for uneven sample sizes, including cases where the sister clade is represented by a single genome^25^.

To first identify groups of isolates with mostly vertically inherited genomes (clonal frame >50%), we applied our mixture model and its associated filters to each of the 176 SGBs that contained more than 5 genomes. Within each SGB, we identified vertically-inherited groups of genomes using the package micropan (bClust, average linkage, ref. ^62^) in R to generate networks of genomes where pairwise vertical inheritance averaged >50%, as determined by our mixture model. In some SGBs, the fraction of recombined genomes gradually decreased with clonal divergence after the initial increase, likely due to the model nearing the limits of its suitable range (Extended Data Fig. 3). Therefore, from each vertically-inherited genome cluster identified, we removed genomes whose average divergence from other cluster members exceeded that of genomes outside the cluster.

We then checked if entire groups of vertically inherited genomes could be putative GWSS clusters. We applied the 5X rule to each vertically-inherited group of genomes within a SGB, by determining if the most closely related isolate outside of the group was more than 5X distant compared to the average clonal divergence within the group. If this condition was met, the entire genome group was considered a putative GWSS cluster. Subsequently, we scanned all vertically-inherited groups of genomes for evidence of GWSSs within the group. Each clade in a maximum likelihood tree, constructed based on whole genome alignments of a vertically-inherited group (phyml, GTR+G+I model, ref. ^63^), was evaluated according to the 5X rule. If the average clonal divergence within a clade was less than 1/5 of that between it and its sister clade, then the clade was identified as a GWSS cluster.

### Validation of putative GWSS clusters in metagenomes

We sought to validate the structure of GWSS clusters as well as the extent of their occurrence in metagenomes representing a large diversity of host conditions and geographic locations. To confirm that the 5X distance gaps for putative isolate GWSS clusters were not due to incomplete or biased sampling, we developed a pipeline that allowed testing of the 5X rule by combining isolate genomes and metagenomes. To this end, we identified a consensus clonal frame (CCF) for each putative GWSS cluster based on isolate genomes and then implemented the 5X rule twice, utilizing two distinct distances: First, the distance of each genome and metagenome sample to the CCF, and second, the pairwise distances between all isolate genome and metagenome samples based on their alignments to the CCF. This procedure is described in detail below.

First, we constructed a database where each isolate-based GWSS cluster is represented by its clonal frame, to ensure that the distances we calculate between metagenomes and each putative GWSS cluster reflect only vertically acquired substitutions. We determined whether the clusters are “nested” (if one cluster completely encompasses another), and only kept the encompassing cluster. For putative GWSS clusters that consist of 3 or more genomes, we extracted the clonal frame of each cluster by removing all recombined fragments in the core genome alignment (Mugsy 1.2.3, ref. ^64^) of the sweep with ClonalFrameML v1.12 (ref. ^65^), and constructed a consensus clonal frame (CCF) by selecting the major allele of each SNP. For GWSSs with only 2 genomes, we extracted the clonal frame by removing all recombined segments (+500bp upstream and downstream) identified by our previous bi-partitioning HMM model, and randomly assigned the clonal frame of one genome as the consensus clonal frame. We then clustered all of the CCFs with fastANI v1.33 (ref. ^52^) and sorted CCFs with ANI >99% into separate databases. This resulted in 6 CCF databases each containing 53-75 clonal frames.

Second, to ensure that the addition of metagenomic samples to the GWSS clusters successfully mitigates potential isolate sampling bias, we acquired a subset of metagenomes that covers many host phenotypes from the curatedMetagenomicData 3.4.2 (ref. ^66^) database. Stool metagenomes were first dereplicated by human subjects, so that for samples sequenced over a time series from the same subject, only the sample with the maximum number of reads was kept. We then grouped the samples by study, age category, disease, and country, and selected up to five metagenomes from each unique group combination. If multiple subjects from the same family were included, we only kept the metagenome for one adult member. This resulted in a collection of 1,477 metagenomes representative of 74 datasets (Table S6).

Third, to ensure that calculation of sequence distance between the metagenomes and the CCF does not represent a mixture of strains, we filtered for metagenomes that were dominated by a single strain from each sweep-containing SGB. Metagenomes were aligned against the MetaPhlAn4 reference database (version Jan. 2022) with MetaPhlAn 4.0.6 (ref. ^23^) default settings, and all polymorphic sites with Phred quality score ≥20 and coverage ≥3 were identified. The allele frequency spectrum was generated for each SGB with >40 polymorphic sites in every metagenome, and SGBs where the 5^th^ percentile of the spectrum exceeded 0.8 was considered as single-strain dominance in that metagenome. This cutoff roughly corresponds to a ratio of 9:1 between major and minor strains.

Fourth, we established a consistent metric to determine the distance of isolate genomes and metagenomes to the CCFs. To determine the distances between single-strain metagenomes and the CCFs, we aligned metagenomes to each of the 6 CCF databases with bowtie 2.5.1 (-X 2000, --no-mixed, --very-sensitive, ref.^67^), and calculated the distances as 1-the popANI metric in instrain 1.7.5 (default settings, ref.^68^). The popANI metric takes into account both major and minor alleles when calculating the distance between metagenomic samples aligned to a reference, making it a population-level measurement of ANI. Furthermore, in order to allow direct comparison between the distances of isolate genomes and metagenomes to CCFs, we converted isolate genomes to synthetic fastq reads (HiSeq 2500 platform, 10X coverage, ART-2016.06.05, ref.^54^), and calculated their distances to the CCFs in the same manner as the metagenomes. We included all isolates that were in the putative sweep clusters as well as up to six isolates that were most closely related to the sweep (“sister isolates”) in the calculation. Eventually, only distances calculated from samples allowing for 2X coverage and 25% coverage breadth (50% for isolate genomes originating from the sweep) were retained for each clonal frame.

Finally, we applied the 5X rule to the sample-to-CCF distances for each GWSS cluster putatively identified based on isolate genomes. All samples, processed as described above, were sorted by their distance to the CCF, from the nearest to the furthest; then, beginning with the third sample in proximity to the CCF, we progressively examined samples in increasing order of distance until identifying a sample whose distance from the CCF exceeded five times the average distance of all samples closer to the CCF. Therefore, a GWSS consisted of least 3 samples. All samples in closer proximity to the CCF than the identified sample are deemed part of the sweep. To avoid identifying GWSSs based on outlier samples that align to the corresponding CCF (i.e., genomes/metagenomes misassigned to a certain SGB), we eliminated all sweeps with fewer than three samples found outside the sweep. Moreover, since some SGBs are known to be mixtures of recently diverged species, and distance gaps can arise from samples far away from the reference CCF not surpassing the coverage threshold, we excluded sweeps if the number of samples within the sweep exceeded 2/3 of the total number of samples, and if, simultaneously, the distance of the closest sample outside of the sweep is more than 1.5 times further from the CCF compared to that determined only by isolates. Finally, in cases where a sample is assigned to two sweeps from the same SGB, we removed the sample from the sweep with less coverage and reran the sweep assignment for the corresponding CCF.

As a final verification of the GWSSs identified by sample-to-CCF distances, we performed an additional 5X test based on pairwise distances. This test included all samples (isolate genomes and metagenomes) in the GWSS cluster as well as up to 6 of the most closely related isolate genomes and metagenomes to the GWSS cluster. For GWSSs occurring in more than 200 samples, we subsampled across the range of distances to the corresponding CCF to obtain a final set of approximately 100 samples for analysis.

Pairwise genetic distances between samples (isolate genomes and metagenomes) were calculated by obtaining the 1-popANI between samples when mapping to the same CCF using inStrain 1.7.5 (ref.^68^). We only considered pairwise samples where more than 25% of the reference CCF was covered by both samples. We then assessed if the average pairwise distance for all samples within the GWSS cluster was less than 1/5 of the average minimal distance of each within-GWSS sample from its closest relative outside of the GWSS cluster. GWSSs that passed this additional 5X test were confirmed as true GWSS clusters.

The phylogenetic structure of all confirmed GWSSs was determined by extracting bac120 marker genes in the GTDB-Tk database (R214, ref. ^43^) from their corresponding CCFs, and inferring a phylogenetic tree from the marker gene alignments, using default settings in GTDB-Tk 2.3.2 (ref. ^43^).

### SGB category assignments

All 176 SGBs where we performed GWSS searches were classified into commensals, pathogens or commensal SGBs that are frequently found in fermented and functional foods (probiotics) according to the following standards. An SGB was categorized as a pathogen if we found that at least a portion of isolate genomes originated from an outbreak by checking the source studies of the isolates. The criterion was that multiple (≥3) genomes were sequenced from the outbreak, since this is also the lowest number of genomes required for the identification of a GWSS. An SGB was designated as a probiotic if literature searches of the corresponding taxa revealed a species commonly found in probiotic products or fermented foods. The remaining SGBs were classified as commensals, and the classification of SGBs can be found in Table S1.

### Calculation of Tajima’s D and dN/dS for GWSS clusters

For calculation of the Tajima’s D of a GWSS cluster, a clonal frame was reconstructed for each isolate and metagenome sample associated with the GWSS, based on the sample’s SNP profile when mapped against the CCF of the GWSS cluster. All reconstructed clonal frames within a GWSS cluster were used to calculate the Tajima’s D of the cluster and its significance level, assuming that D follows a beta distribution using the tajima.test function in the pegas package (ref.^69^) in R. Genes under positive selection in each GWSS cluster were classified as those that had sufficient coverage (coverage depth >2X, coverage breadth >50%) and dN/dS >2 compared to their counterparts outside of the cluster. The choice of 2 as the dN/dS cutoff was based on the 95% confidence interval for the dN/dS of genes in closely related genomes evolving under neutral processes^28^. To ensure that the selection pressure was on the gene in the GWSS cluster and not its counterpart outside the cluster, we required the gene to be under positive selection when compared to least two out of the three of the most closely related sister genomes or metagenomes of the GWSS. The dN/dS of genes in each sample was calculated using the SNP information when mapped against genes extracted (prodigal, ref.^70^) from the CCF of the GWSS cluster using inStrain 1.7.5 (ref. ^68^).

### GWSS cluster age estimation

The age of each GWSS cluster was calculated using two independent methods by (i) dividing the maximum pairwise SNP distances within each cluster by 2, and then with a constant molecular clock of 1-10 mutations genome^−1^ yr^−1^, and (ii) estimating a molecular clock from strains in twin metagenomes or metagenomic time series. Within each GWSS cluster, the SNP distances between strains in two samples were calculated by normalizing the population_SNPs metric (inStrain 1.7.5, ref. ^68^, defined as sites where coverage is >5X with no shared alleles between the samples) by the percent of reference CCF with >5X coverage in both samples. This pairwise SNP distance calculation was performed exclusively on samples where more than 25% of the reference CCF had >5X coverage.

We were able to estimate the metagenomic molecular clock of 9 SGBs from the metagenome time series or twin metagenomes, by finding all strains that persisted in individuals over a period of time or were shared between twins. We retrieved all metagenomes from the same human subject that were at least sampled one year apart from curatedMetagenomicData 3.4.2 (ref.^66^). If multiple time points were sampled for the same human subject, we selected the two time points that were furthest apart. We also retrieved all metagenomes and their related metadata from 250 adult twins from the TwinsUK study^71^. Twins were assumed to have identical strains when they were living in the same household, and the years that the twins had lived apart were assumed as the time the strains had to accumulate mutations. The genetic distance between strains in each metagenome and the CCF of their corresponding GWSS were calculated in the same manner as in the previous section (“Validation of putative GWSS clusters in metagenomes”). To account for shifts in strain dominance and strain replacement events over time, we only considered metagenomes from the same person/twin pair to be sharing strains from the same GWSS, if one metagenome was more closely related to the reference CCF of the GWSS compared to the threshold previously used to establish the GWSS cluster, while the other metagenome was closer to the CCF than half of any other metagenomes not included in the GWSS cluster.

For each SGB, the metagenomic molecular clock was expressed as a linear function, with the SNP distance between shared strains in metagenomes as the independent variable, and the time difference between the metagenomes as the dependent variable. When SGBs contained shared strains in two or more metagenome pairs, the linear function was determined as the best fit across all data points. In cases where the SGB only had shared strains in a single metagenome pair, the function was a line passing through the single data point and the origin. We eventually estimated the age of every GWSS cluster belonging to the 9 SGBs by extrapolating the corresponding metagenomic molecular clock to the maximum pairwise SNP distance of the GWSS cluster.

### Curve fitting for measuring recombination rates

Since in most SGBs the fraction of the genome that has undergone recombination increased linearly as the number of mutations in the clonal region increased, and subsequently plateaued, the slope of the linear segment of the recombined fraction-mutation plots is a measure of recombination rate. We thus segmented all recombined fraction-mutation plots (Extended Data Fig. 6) for commensal bacteria to find their linearly increasing regions, using the R package dpseg 0.1.1 (ref. ^72^). Since sometimes a large number of data points clustered at low divergence and this could lead to over-segmentation of the scatter plot, we subsampled the plot to 100 data points when there were >100 data points with less than 2,000 mutations. A total of 4 parameter combinations that included breakpoint penalty, minimal and maximal segment length were tested for the curve fitting [(0.2,20,40), (0.1,10,20), (0.1,5,10), (0.2,20, all data points)]. If the first linear fragment of the fit had R^2^>0.8, then this fragment was determined as the linearly increasing region. If R^2^>0.8 was satisfied under multiple parameter combinations, then the combination that had the maximum R^2^ or allowed all data points to fit to a single linear fragment with R^2^>0.8 was used. Otherwise, consecutive linear fragments with similar (within 75%) slopes were combined and refit as one fragment, and the first fragment with R^2^>0.33 was determined as the linearly increasing region. For SGBs where automated segmentation was not satisfactory, we manually identified the linear range of increase. Finally, for all the identified linearly increasing regions of each SGB, we added a point (0,0) to the data points in the region, and applied a linear regression model passing through the origin. The slope of the linear regression model was used as the recombination rate, and we were able to measure the recombination rates for 45/46 of the commensal SGBs with sweeps, and 52/95 of those without sweeps. The lower fraction of satisfactory fits in the SGBs with no confirmed sweeps was due to both fewer genomes per SGB, and a more frequent lack of data points in the linearly increasing fraction of the recombined fraction-mutation plots. All curve fittings are shown in Extended Data Fig. 6, and all measured recombination rates are in Table S7.

### GWSS identification from StrainPhlAn marker gene trees

Because we were interested in testing associations of GWSS clusters with human disease or physiological states, we explored the feasibility of identifying GWSSs based on phylogenetic distances of marker genes extracted from the metagenomes since this approach can be more easily scaled up to large metagenomic datasets. Since our SGB classifications were based on the MetaPhlAn4 database, we performed strain-level marker gene profiling for SGBs in metagenomes with StrainPhlAn4, and tested how and to what extent the 5X rule could be extended to StrainPhlAn4 marker gene trees to identify GWSS clusters. We set up two criteria for calling a GWSS cluster from the marker gene tree: (i) The normalized average genetic distance within a marker gene based GWSS cluster needs to be smaller than a normalized cutoff based on previously identified GWSS clusters, and (ii) the phylogenetic distance between the proposed GWSS clade and its sister clade exceeds five times the average distance within the GWSS clade.

To define the cutoff for the first criterion, we constructed mock metagenomes for all isolate genomes within GWSS clusters. Each mock metagenome for an isolate genome consists of synthetic fastq reads for the target genome at 20X coverage (ART-2016.06.05, -ss HS25 -f 20, ref.^54^), and a randomly selected isolate genome from every other GWSS-containing SGB at 1X coverage. Therefore, the total number of mock metagenomes for each SGB is the number of isolate genomes identified within GWSS clusters for that SGB. For each SGB, strain-profiling was performed for each mock metagenome with StrainPhlAn4 against the MetaPhlAn4 reference database (version Jan. 2022, ref.^23^), resulting in a tree that was built using marker genes from all the isolate-based mock metagenomes and marker genes extracted directly from all other isolate genomes in the SGB. The cutoff was set as SGB-specific normalized phylogenetic distance (nGD) thresholds that optimally separated isolate pairs within GWSS clusters from isolate pairs that had only one isolate genome in the GWSS cluster. nGDs were calculated as leaf-to-leaf branch lengths on the SGB marker gene tree normalized by their median. For SGBs with at least 50 pairs of isolates within the GWSS cluster, nGD cutoff thresholds were defined based on the value that would maximize the Youden’s index (R package cutpointr, ref.^73^), unless the value exceeded the 5^th^ percentile of the isolate pairs which had only one isolate genome in the GWSS cluster. For SGBs with less than 50 total within-GWSS isolate pairs, the nGD corresponding to the 3^rd^ percentile of the isolate pairs with only one isolate genome in the GWSS cluster was used as the cutoff.

StrainPhlAn marker gene trees were constructed for each SGB with the same isolate and metagenome samples previously used to identify GWSSs. For 32/46 of the commensal bacteria, at least 2/3 of the samples in previously identified GWSS clusters are retained in GWSS clusters called from the StrainPhlAn marker gene trees (Extended Data Fig. 7a). Also, since GWSS clusters identified from marker gene trees are not necessarily the expansion of isolate based GWSS clusters but can be purely metagenome based, for the majority of the SGBs (33/46), GWSS clusters called from the StrainPhlAn marker gene trees include more samples than those included in the previously identified GWSS clusters (Extended Data Fig. 7b).

### Association studies of GWSS clusters in metagenomes

Associations between GWSS clusters and two human metrics, colorectal cancer (CRC) and seniority, were examined for each SGB. The two metrics were chosen because they were most likely to have sufficient numbers of samples spanning multiple biogeographies. To start, we assembled a metagenome database comprising 17 datasets, which collectively included 3,941 metagenome samples sourced from curatedMetagenomicData 3.4.2 (ref.^66^). Among these datasets, 6 specifically contained samples from colorectal cancer (CRC) patients, constituting the entirety of CRC datasets available in the database (Table S8). For all 46 commensal SGBs with previously confirmed GWSSs, strain-profiling was performed with StrainPhlAn4 against the MetaPhlAn4 reference database (version Jan. 2022, ref.^23^).

Markers for each SGB were extracted from all isolate genomes and all metagenomes with single-strain dominance for the SGB (see section “Validation of putative GWSS clusters in metagenomes”). For each SGB, all extracted markers were aligned, filtered and constructed into a maximum likelihood tree according to the default settings under the accurate mode of StrainPhlAn4. The two criteria for identifying GWSS clusters in StrainPhlAn marker gene trees were applied to each SGB. A total of 1,063 GWSS clusters were identified in 40 of the 46 commensal SGBs examined.

As further preparation for the association analysis, we performed some additional filtering and metadata curation for all the samples involved in each SGB marker tree. Since age/CRC information for isolate genomes was often not available, we removed all isolate genomes from the SGB trees. For samples originating from the same subject or subjects in the same family, we only kept one sample at random from each subject/family for each SGB tree. We also removed all samples in the “newborn”, “child”, “school-age” age category. A total of 693 GWSS clusters remained in 36 of the 46 commensal SGBs examined after this additional filtering. Finally, we classified samples from subjects aged ≥65 years old as “seniors”, and the remaining as “adults”. Samples labelled as “carcinoma” or “CRC” were classified as “CRC”, and the remaining as “non-CRC”.

Associations between GWSS clusters in each SGB and CRC/seniority were examined by building a general linear model with stepwise, forward variable selection and FDR correction. We asked whether being in a certain sweep or not has a positive or negative impact on the sample being from CRC patients or a senior with the formula Y (CRC or senior) = ß_1_S_1_+ß_2_S_2_+…ß_n_S_n_+μ, where S_1_, S_2_,…S_n_ represent GWSS clusters detected in each SGB. The forward selection was performed with the R package SignifReg (v4.3 ref. ^74^), under the criteria that a new predictor is added to the model if the addition of the predictor further minimizes the model p-value, and every individual predictor remains significant at p_adj_<0.05 after FDR correction. All selected sweeps were further tested for geographical biases to ask if sweeps are dominated by samples from certain countries by performing a chi-squared test for the country distribution in each sweep. Country-specific sweeps were identified with the same procedure but only using samples that originated from 7 selected countries where CRC samples were available (JPN, IND, CHN, FRA, ESP, DNK, USA).

Associations between SGBs and CRC/seniority were performed in the same manner as for GWSS clusters within each SGB, with the formula Y (CRC or senior) = ß_1_SGB_1_+ß_2_SGB_2_+…ß_n_SGB_n_+μ, where SGB_1_, SGB_2_,…SGB_n_ represent individual SGBs. Due to the large number of metagenomes associated with each SGB, we did not test for geographical bias within each SGB.

### Identification of sweep-specific genes

To identify gene clusters (i.e., consecutive genes likely belonging to the same operon) that were specific to each GWSS cluster from *Bacteroides fragilis* (SGB1855), we first predicted all protein-coding genes in the CCF and isolate genomes (Prodigal v2.6.3, ref. ^70^) from each of the 6 isolate-based sweep clusters.

The protein-coding genes in each CCF were then pairwise aligned at the protein and nucleotide level (Blast v2.15.0+, ref. ^75^), and proteins found in a single CCF were selected as sweep-specific genes. Specifically, the selected proteins had no homolog in the other 5 CCFs after filtering for alignments with over 60% amino acid identity and alignment length. We then required the genes encoding the selected proteins in each CCF to be identical at the nucleotide level in isolate genomes in the corresponding sweep cluster. Selected protein-coding genes occurring sequentially in the corresponding CCF in sets of 3 or more were designated sweep-specific gene clusters, which were annotated using EggNOG (emapper v.2.1.12, database v.5.0.2, ref. ^76^) and Prokka (v. 1.14.6, ref. ^77^). Lastly, to test for Clusters of Orthologous Groups (COG) categories or Pfams enriched in the sweep-specific gene clusters, annotations in the gene clusters were compared to those in the entire CCFs using a Fisher’s exact test with Bonferroni correction.

### Statistical analysis

Statistical analyses and graphical representations were performed in R (v.4.2.1, ref. ^78^) using base R statistical functions and ggplot2 (v3.5.1, ref. ^79^), ggpubr (v0.6.0,ref. ^80^), ggtree (v3.4.4, ref. ^81^), ggtreeExtra (v1.6.1, ref. ^82^), and ComplexHeatmap (v2.12.1, ref. ^83^). Correction for multiple testing (Benjamini– Hochberg procedure) was applied when appropriate and significance was defined at p_adj_ < 0.05. All tests were two-sided. To access differences between two groups, the student t-test was performed on data that passed the Shapiro-Wilk normality test; otherwise, a Wilcoxon rank sum test was performed. Correlations were assessed with Spearman’s tests. All geographical biases in the datasets were accessed either with a chi-squared test or a Fisher’s exact test.

### Ethical compliance

For the Austrian isolate collection, study approval was granted by the ethics committee of the Medical University of Vienna (EK-Nr: 1617/2014, 1910/2019). All study participants gave written informed consent before study inclusion. The study was conducted in accordance with the ethical principles of the Declaration of Helsinki. The analysis of the Global Microbiome Conservancy isolate dataset was conducted with authorized access to data from the database of Genotypes and Phenotypes (accession phs002235.v1.p1), under approval from the US National Human Genome Research Institute.

## Data and Code availability

The PopCoGenomeS source code is available at https://github.com/cusoiv/PopCoGenomeS. All other code is available upon request. All newly sequenced genomes are uploaded to NCBI under the BioProject PRJNA1101861, and accession numbers for all isolate and metagenomes used are available in Tables S5, S6 and S8.

## Notes

### Competing Interest Statement

The authors have declared no competing interest.

