## Supplementary material for "Genome-wide sweeps create fundamental ecological units in the human gut microbiome": All Supplementary Figures

#### Extended Data Figure 1

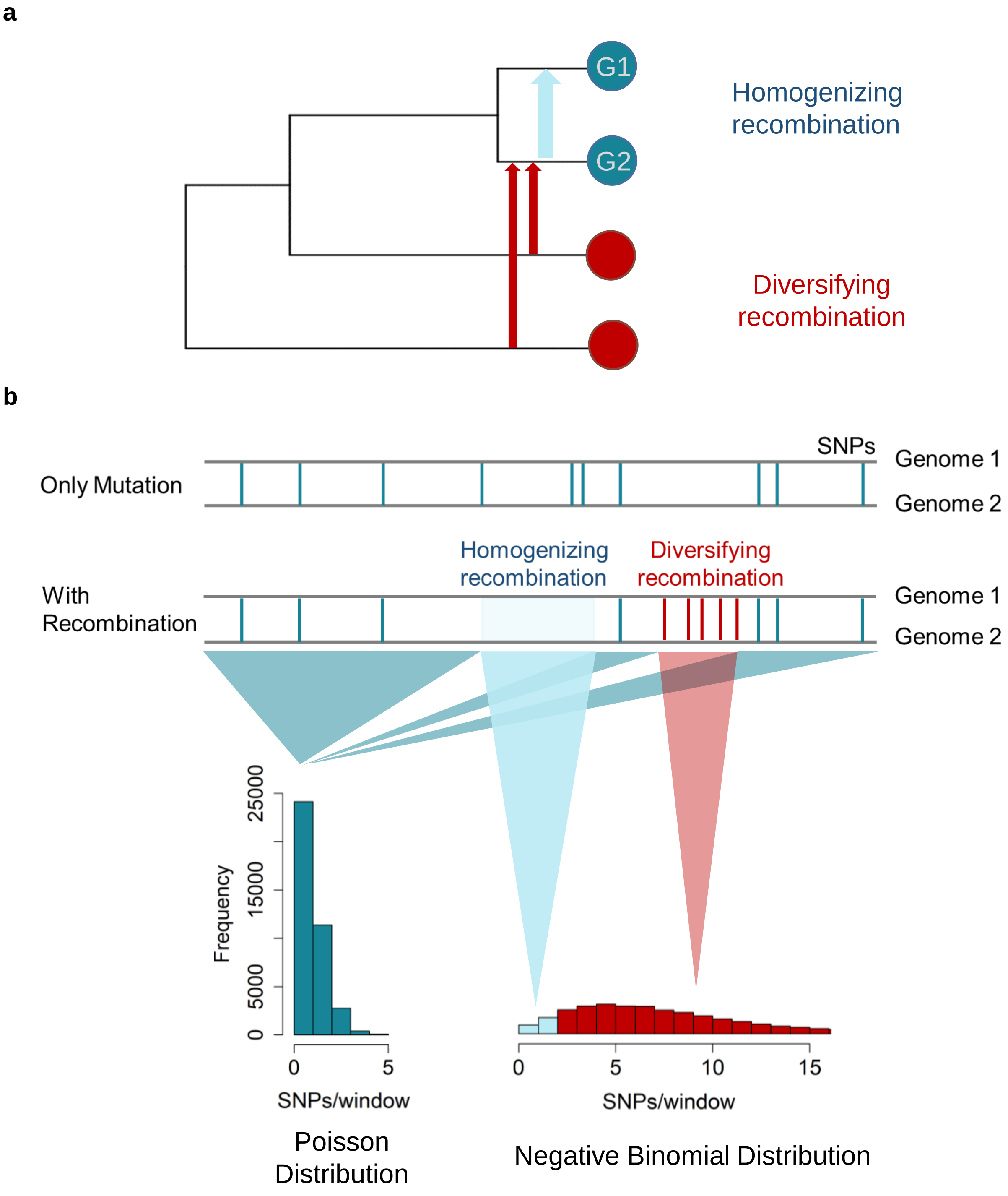


**Extended Data Fig. 1. Single nucleotide polymorphism patterns across pairwise genome alignments reflect point mutations and recombination. a.** Illustration of two types of recombination. For the focal pair of genomes (G1 and G2, dark cyan), recombination can happen between the two genomes (homogenizing recombination, light blue), or with phylogenetically more distant genomes (diversifying recombination, red). **b.** Recombination alters the SNP distribution between two genomes by either eliminating SNPs (homogenizing recombination, light blue) or by increasing the SNP density (diversifying recombination, red). The distribution of the number of SNPs in sliding windows across pairwise genome alignments can be expressed as the fractional sum of a Poisson distribution (point mutations), and a Negative Binomial distribution that accounts for both diversifying and homogenizing recombination.

**Extended Data Figure 2**

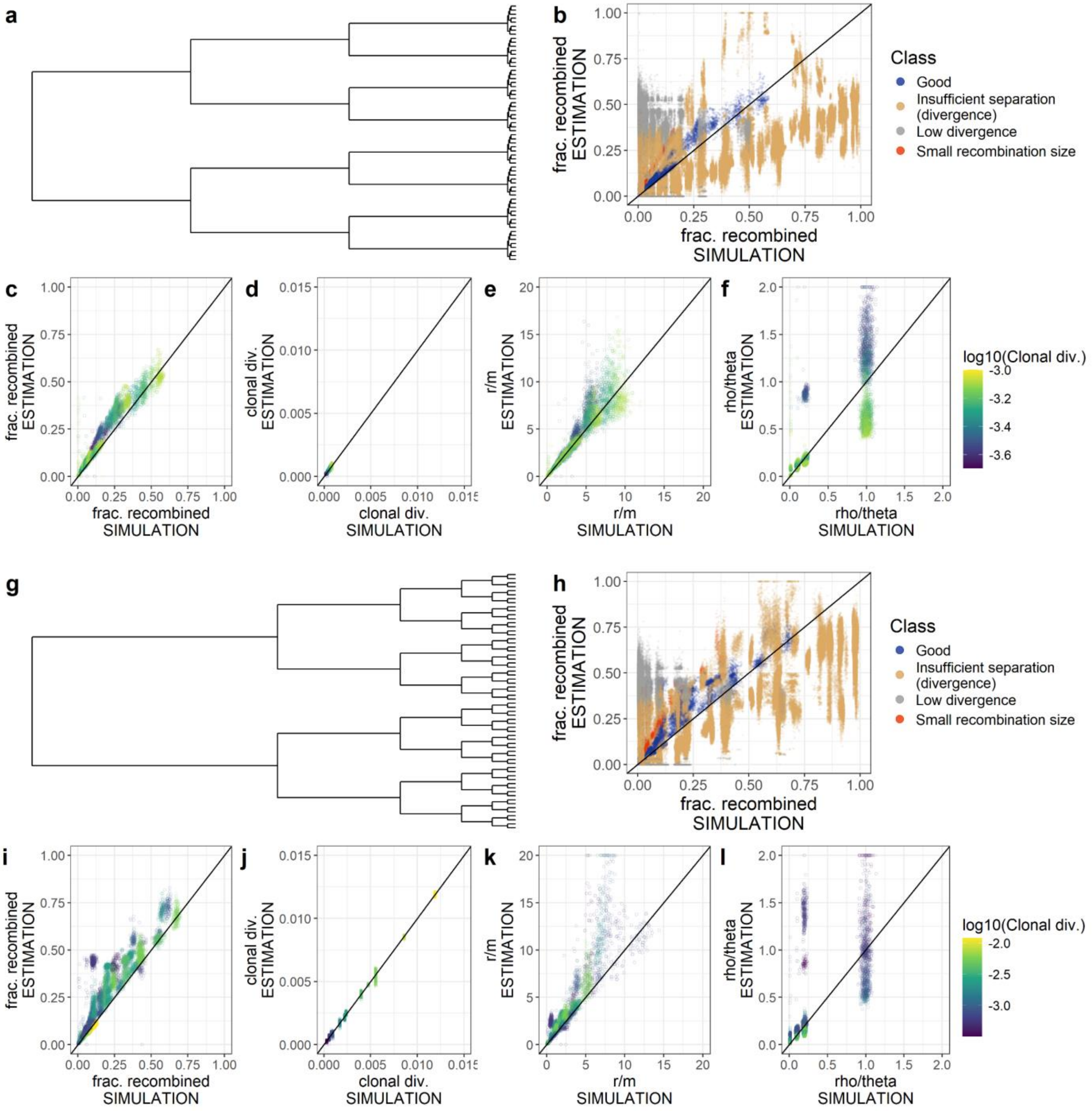


**Extended Data Fig. 2. Validation of the mixture model for recombination estimation with simulated data. a.** Phylogenetic structure of sets of 64 simulated genomes evolved along a phylogenetic tree with sweep clusters under 144 parameter combinations (Methods). **b.** Comparison of the recombined genome fraction in the simulation and estimated by the model. Different colors represent categories of genome pairs where the model accurately estimates the recombined genome fractions (dark blue), or is insufficient due to different reasons: yellow, insufficient differentiation of the clonal and recombined fragments due to similar divergences (<2.5-fold difference); grey, overall divergence too low (<0.0075%); red, insufficient length of recombination fragments (exponential distribution with a mean of 200 base pairs). **c-f.** Comparison between values in the simulation vs. estimated values within the functional range of the model for recombined genome fraction, clonal divergence, the effective recombination rate r/m, and relative recombination to mutation occurrence rate rho/theta. **g-i.** same as a-f but for sets of 64 simulated genomes with balanced phylogenetic tree structure.

### Extended Data Figure 3

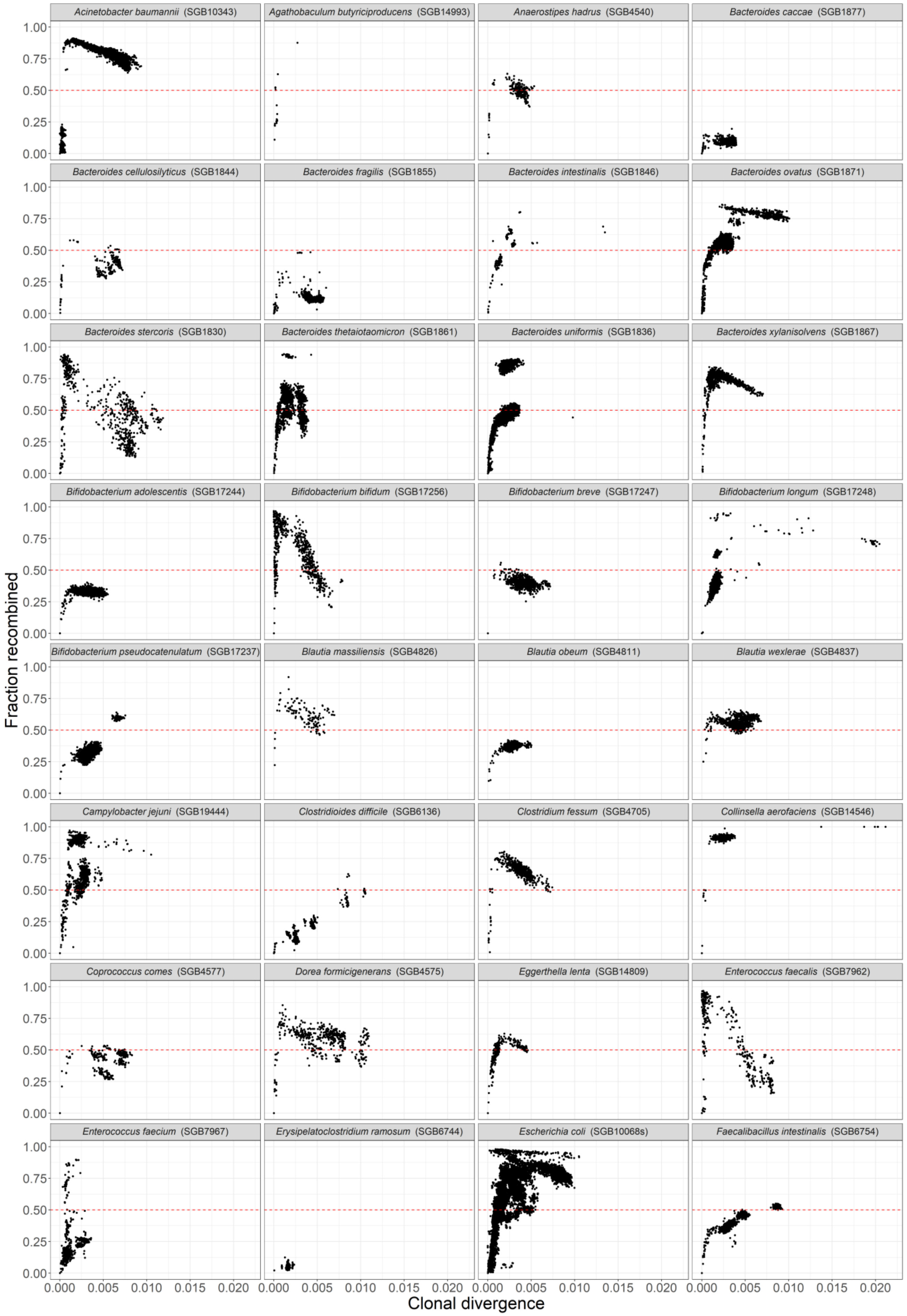

**Extended Data Figure 3 (Continued)**

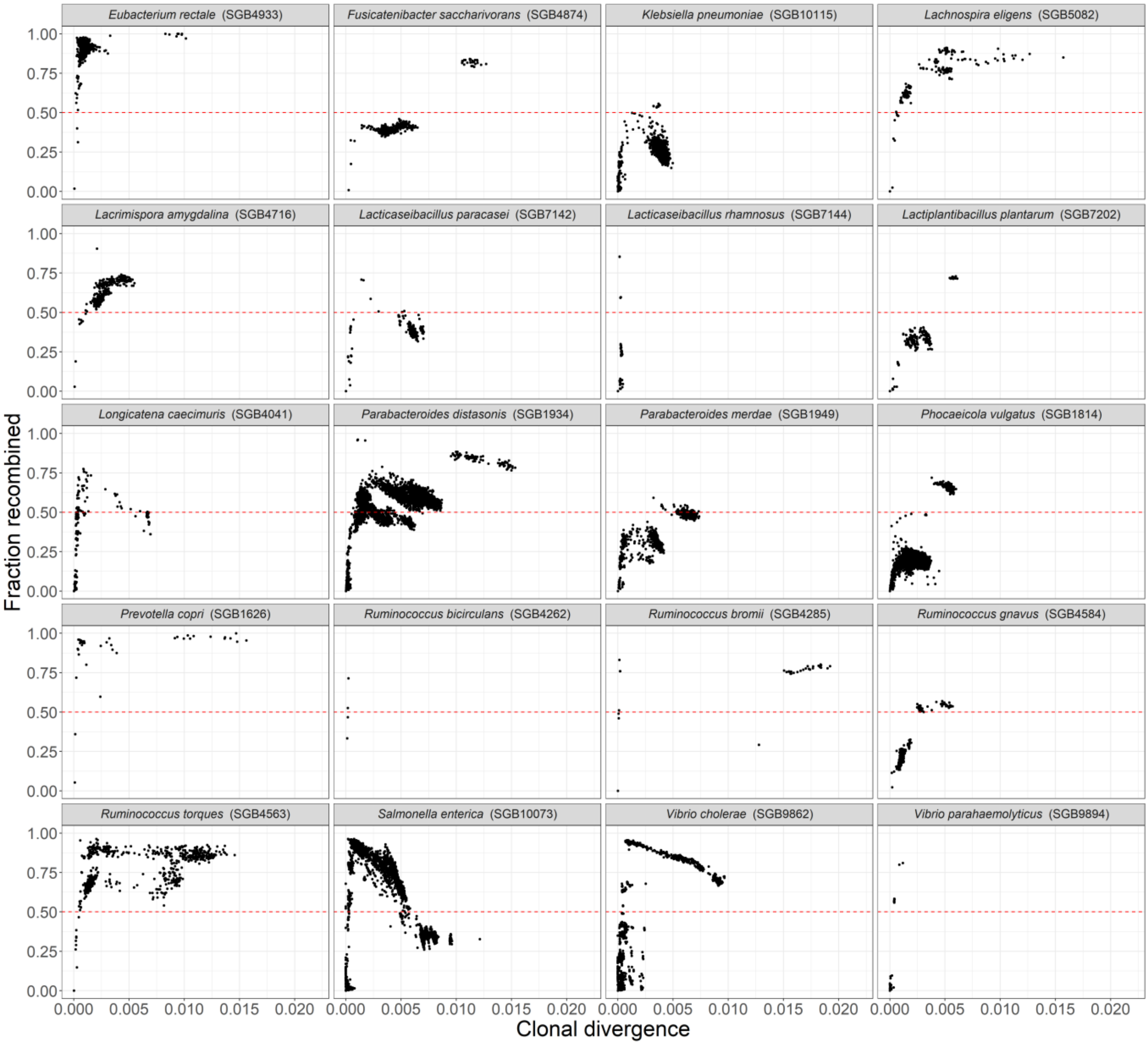


**Extended Data Fig. 3. Recombination-divergence relationships among isolate genomes.** The relationship between clonal divergence and fraction of the genome that has undergone recombination for all SGBs with more than 20 isolate genomes (52 out the total 176 SGBs), after implementation of filtering and corrective measures. Each dot represents a pair of isolate genomes and the red line at 0.5 is the cutoff chosen for predominantly vertical inheritance (<50% recombined). Clusters of genomes below this threshold were subsequently utilized for downstream GWSS searches.

**Extended Data Figure 4**

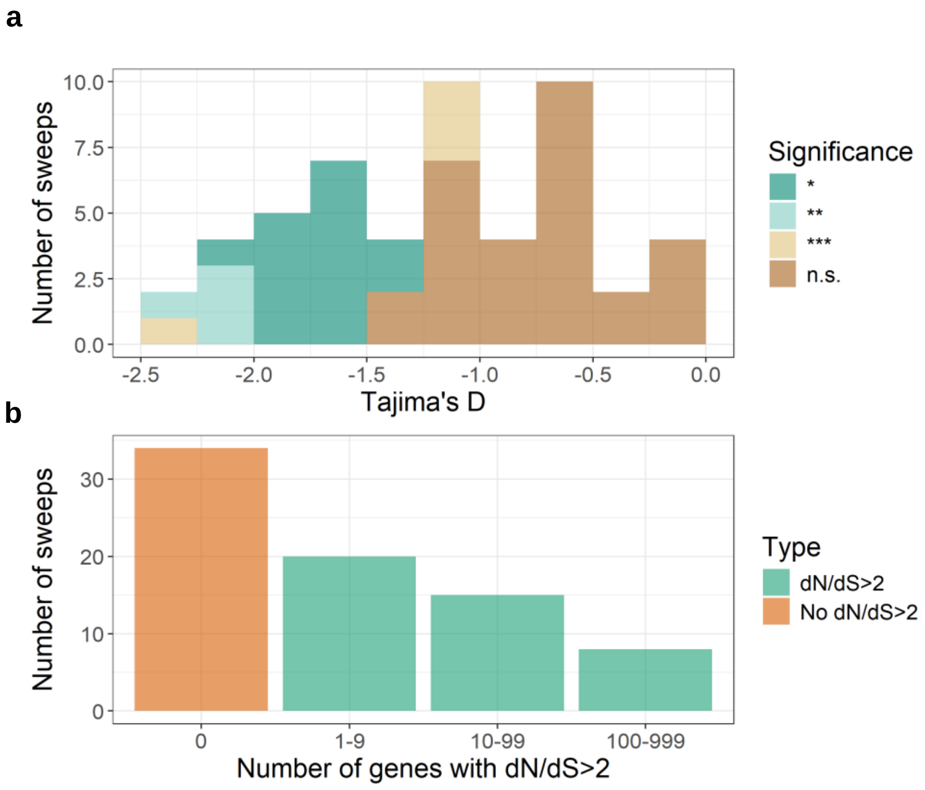


**Extended Data Fig. 4. Population genomic signatures of genome-wide selective sweep clusters. a.** Histogram of the Tajima's D for all GWSS clusters detected in commensal gut bacteria. ***, p<0.001; **, p<0.01, *, p<0.1. **b.** Histogram of the number of genes with dN/dS >2 in the consensus clonal frames of all GWSS clusters for commensal gut bacteria.

**Extended Data Figure 5**

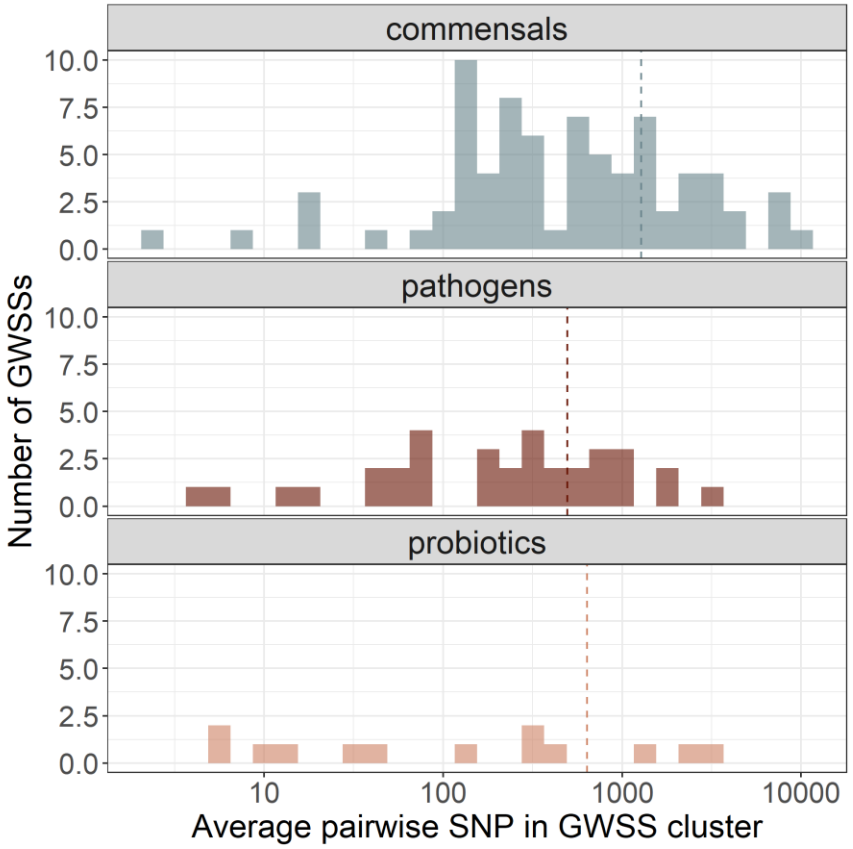


**Extended Data Fig. 5. Average SNP distance within genome-wide selective sweep clusters for commensal, pathogenic and probiotic bacteria.** Dashed line in each histogram represents its mean. See Methods for how SGBs are assigned to the three different categories**.**

#### Extended Data Figure 6

**a**

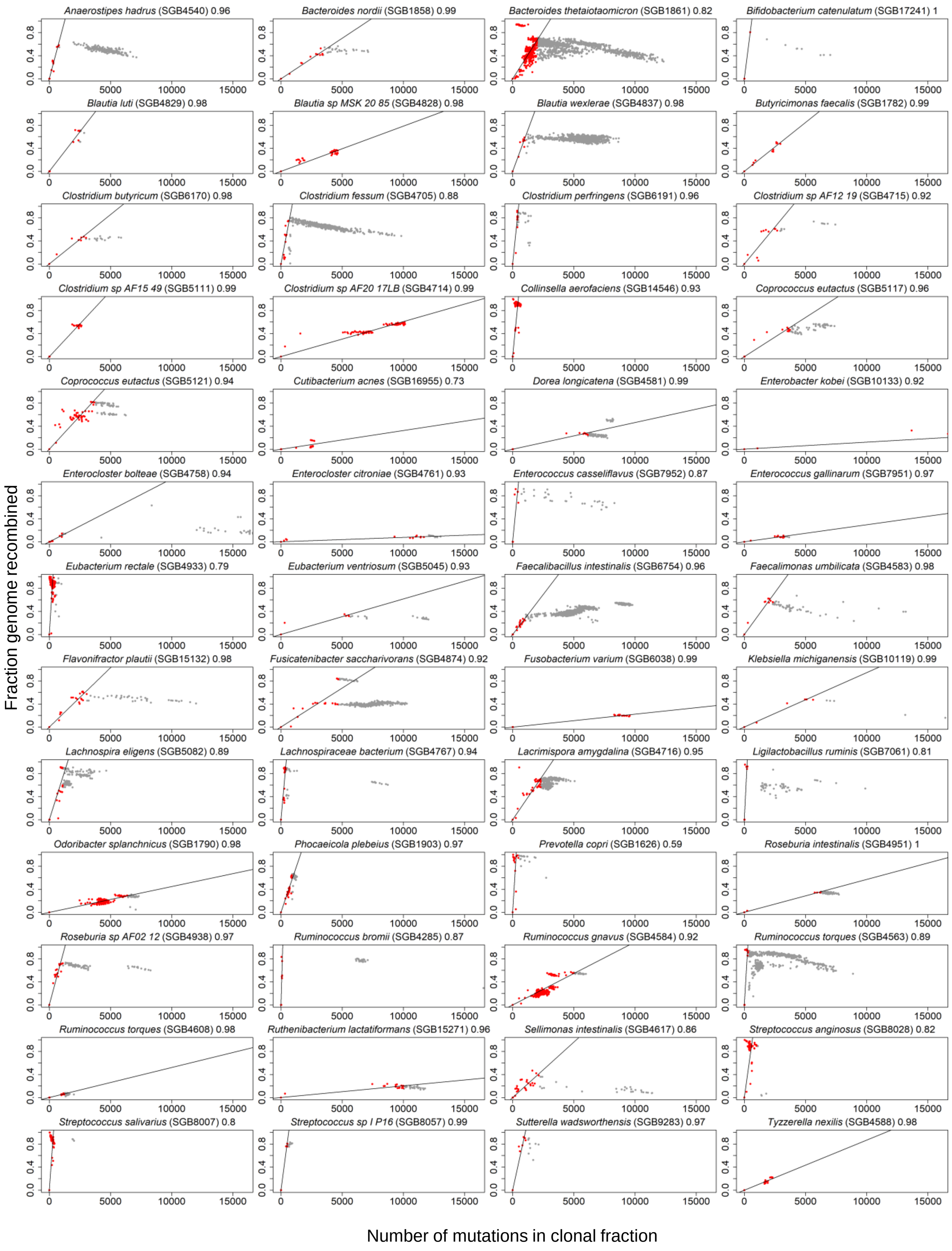

**Extended Data Figure 6 (Continued)**

**b**

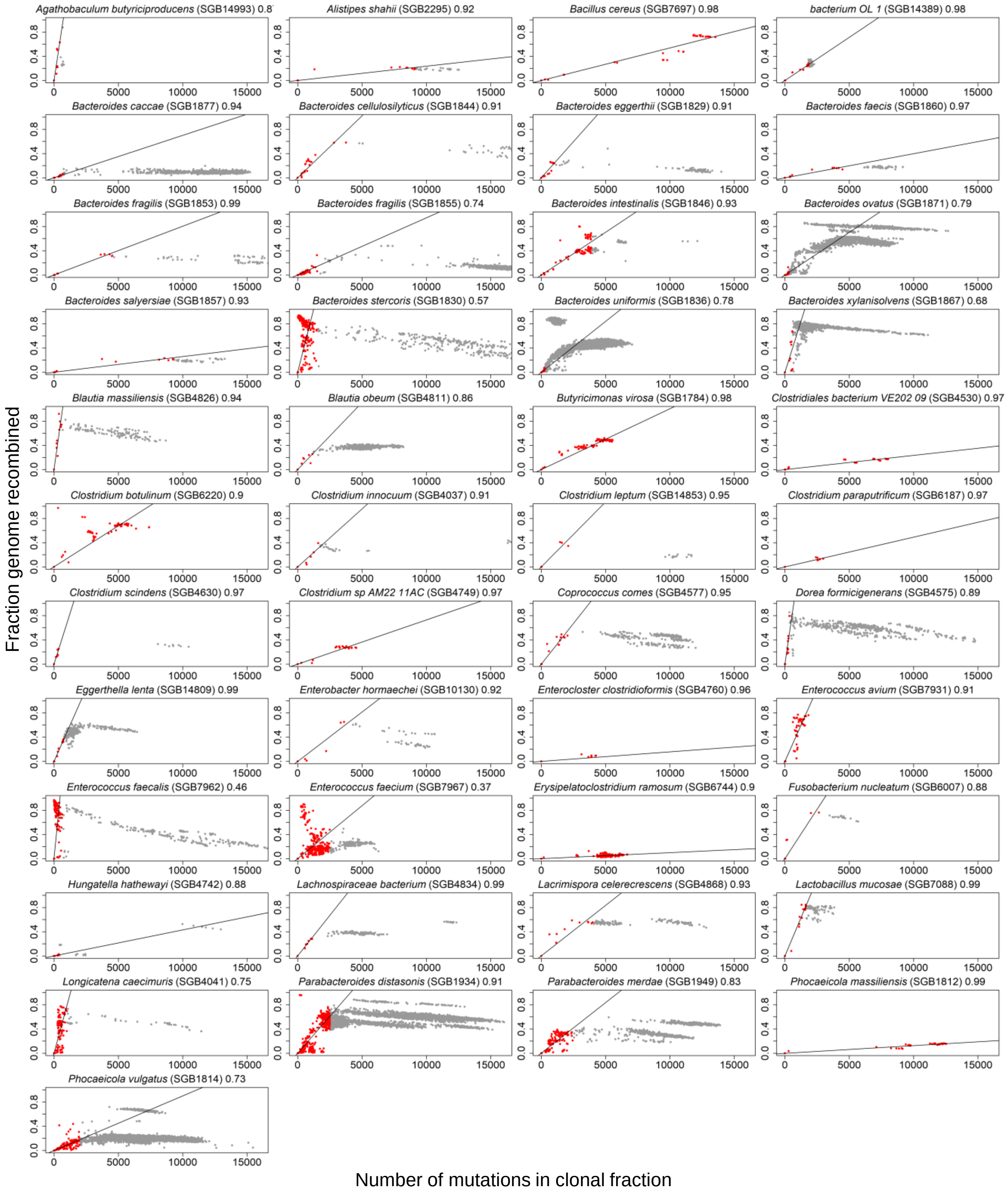


**Extended Data Fig. 6. Recombination rates vary among SGBs.** Curve fitting for all **a.** SGBs with no confirmed GWSSs and **b.** SGBs containing GWSSs with satisfactory fits ($R^2$>0.33) for the fraction of recombined genome vs. number of mutations in the clonal fraction. Each dot represents a pair of genomes compared, with red dots indicating those used for curve fitting, and the black line being the fitted line. The number following the alphanumerical SGB names in the title of each plot represents the $R^2$ of the fit.

#### Extended Data Figure 7

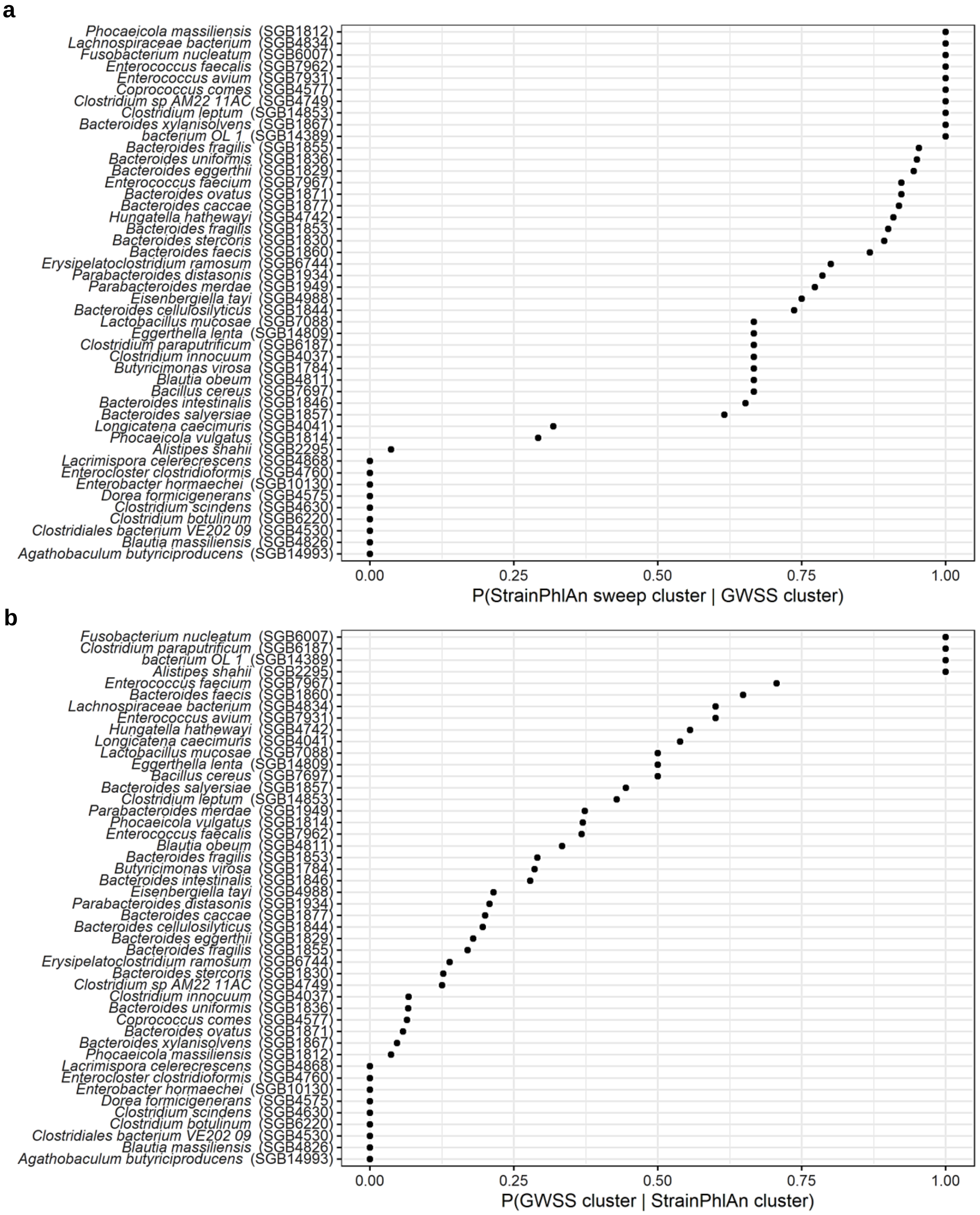


**Extended Data Fig. 7. Comparison of genome-wide sweep clusters identified using isolate-based clonal frames and StrainPhlAn marker genes for each of the 46 commensal SGBs. a.** The probability that samples (including isolate genomes and metagenomes) within GWSS clusters identified using isolate-based clonal frames also fall within sweep clusters called from the StrainPhlAn marker gene trees. **b.** The probability that samples (including isolate genomes and metagenomes) within sweep clusters called from the StrainPhlAn marker gene trees, also fall within GWSS clusters identified using isolate-based clonal frames.

#### Extended Data Figure 8

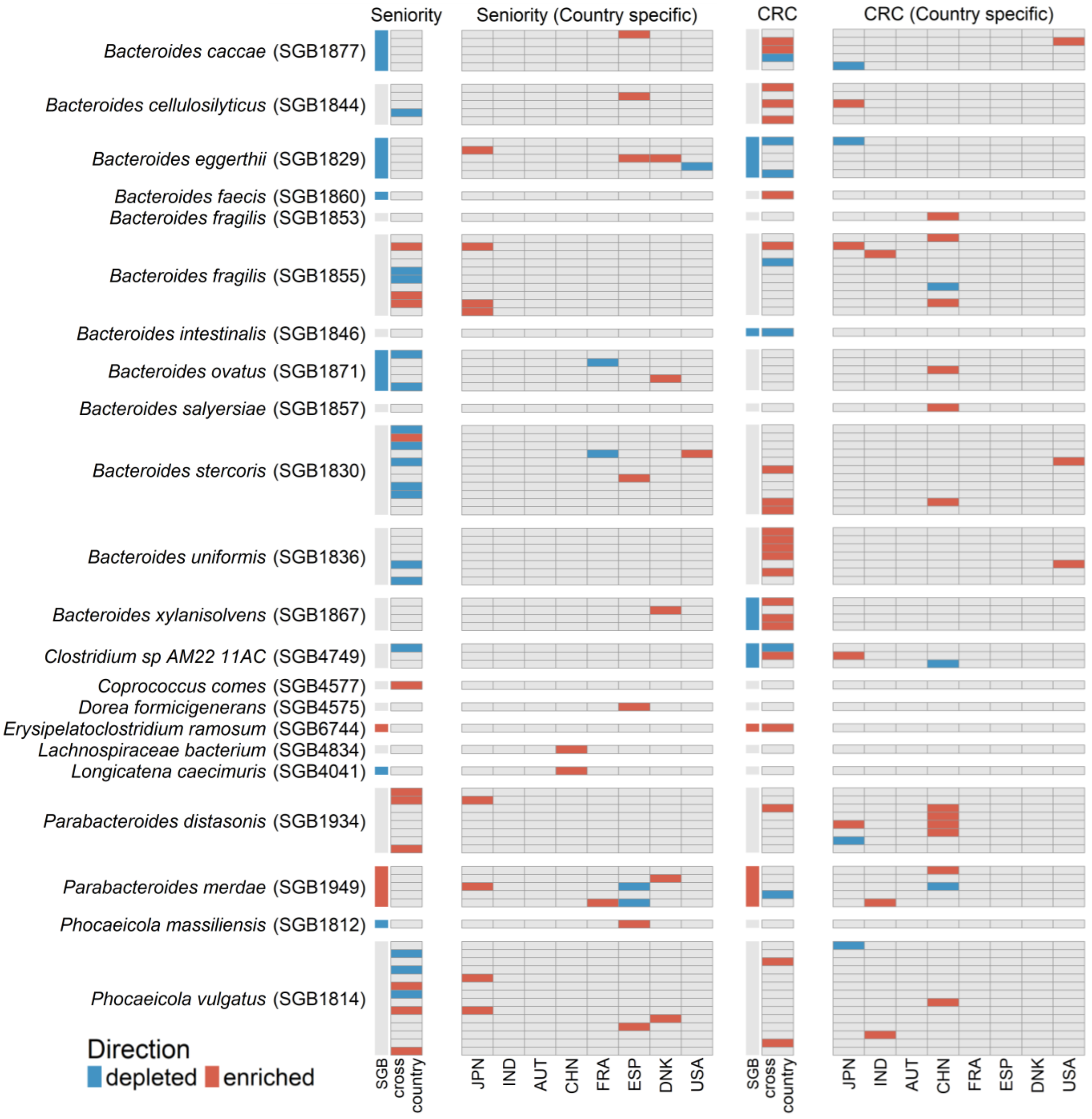


**Extended Data Fig. 8. Genome-wide selective sweep clusters and SGBs associated with host conditions.** Heatmaps of statistically significant ($p_{adj}<0.05$) associations between GWSS clusters and SGBs and seniority/CRC. For both seniority (left two panels) and CRC (right two panels), each row represents a GWSS cluster and the rows are split by the SGBs that the GWSS clusters belong to. For both conditions, there is one panel (left) that represents associations that span multiple datasets and show no significant geographical bias ($p>0.05$, chi-squared test), and another panel (right) that shows significant associations ($p_{adj}<0.05$) that can only be identified in specific countries. Vertical bars on the side of each panel represent whether the SGB that the GWSS clusters belong to is significantly associated with seniority or CRC ($p_{adj}<0.05$).

### Extended Data Figure 9

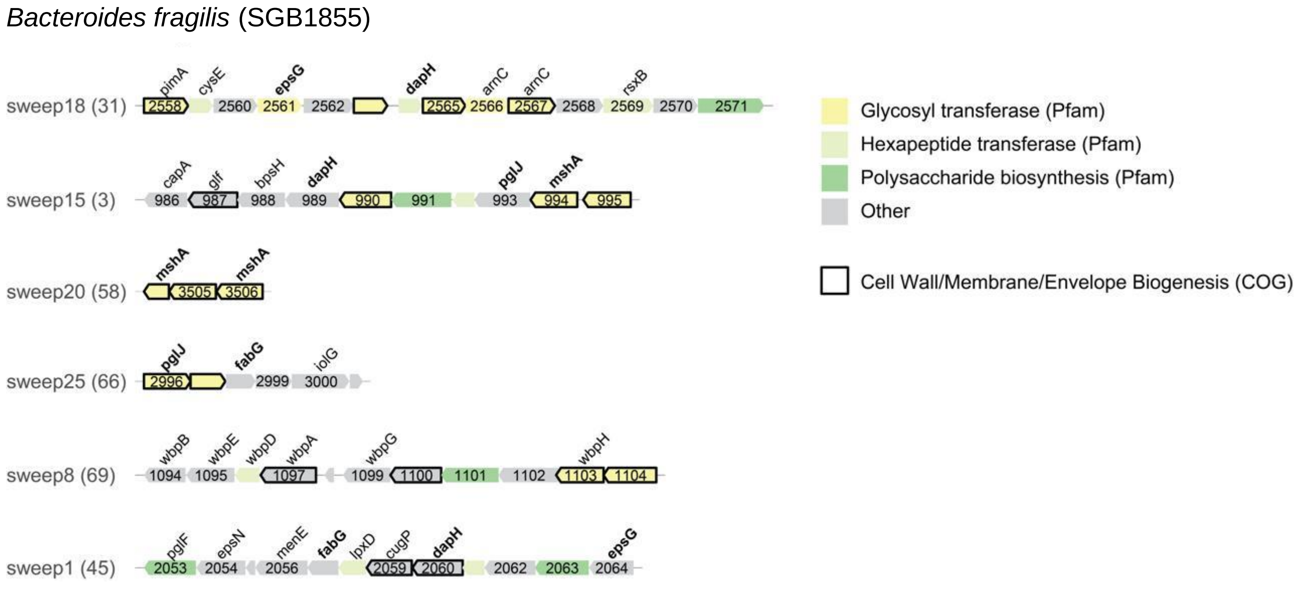


**Extended Data Fig. 9. Gene clusters unique to the clonal frames of the genome-wide sweep clusters of *Bacteroides fragilis*.** Predicted proteins within the clonal frame of 6 sets of isolate genomes belonging to GWSS clusters within *B. fragilis* (SGB1855) were compared to each other. Three or more sequential predicted proteins in a clonal frame from a GWSS were selected if they were missing from all other clonal frames of the GWSS clusters. The encoding genes were further required to be identical in isolate genomes belonging to the GWSS. The resulting gene clusters were then tested for overall enrichment of annotated COG categories and pfams (Fisher's exact test with Bonferroni correction). The COG category M (black border) and pfams associated with glycosyl transferases, hexapeptide transferase, and polysaccharide biosynthesis (colored) were significantly enriched (adj. p-value $< 5\times10^{-6}$). An example gene cluster (cluster id in parentheses after sweep name) is shown for each GWSS cluster based on the highest number of non-hypothetical annotations, while the annotations for all gene clusters are presented in Table S3. The number on each gene represents its ORF number within the corresponding consensus clonal frame, and where available, a putative gene name is provided; those appearing multiple times across the different GWSS clusters are bolded.
